# FAIRyMAGs - a series of FAIR Galaxy workflows for the generation of metagenome assembled genomes

**DOI:** 10.64898/2026.07.31.741430

**Authors:** Paul Zierep, Mina Hojat Ansari, Patrick Bühler, Santino Faack, Bruno Fosso, Giuseppe Defazio, Martin Beracochea, Santiago Sanchez, Alexandra Hottmann, Bérénice Batut

## Abstract

Advances in whole-genome sequencing (WGS) technologies have enabled large-scale recovery of metagenome-assembled genomes (MAGs), providing unprecedented insights into microbial diversity across diverse environments. However, the reconstruction of MAGs remains computationally demanding and methodologically complex, requiring the integration of multiple tools for quality control, assembly, binning, refinement, and annotation. Existing workflows often rely on scripting-based implementations, constrain user-driven modification and stepwise execution, and require advanced expertise in high-performance computing (HPC) system administration, thereby limiting accessibility, reproducibility, and adaptability.

Here, we present FAIRyMAGs, a Findable, Accessible, Interoperable, and Reusable (FAIR)-compliant, modular pipeline implemented within the Galaxy platform for the generation and analysis of MAGs. FAIRyMAGs consists of six interconnected workflows covering all major steps of MAG reconstruction, including read preprocessing, host and contaminant removal, assembly, binning, dereplication, and downstream taxonomic and functional annotation. The workflows are accompanied by extensive training material, including tutorials, a learning pathway, FAQs, test datasets and video walk-throughs by domain experts, supporting community adaptation.

By leveraging Galaxy’s graphical interface and federated infrastructure, FAIRyMAGs enables users to execute complex analyses on public or private compute resources without requiring local installation or workflow programming expertise. The modular design further supports flexible adaptation, iterative optimization, and seamless integration of new tools contributed by the community.

To demonstrate applicability, FAIRyMAGs was applied to four real-world microbiome datasets spanning various host-associated and environmental systems. These analyses revealed substantial variability in MAG recovery, community complexity, and clustering structure, underscoring the importance of flexible workflows adaptable to dataset-specific characteristics. Overall, FAIRyMAGs provides an accessible, extensible, and reproducible framework for genome-resolved metagenomics, reducing technical barriers and enabling methodological innovation through community-driven development within the adaptable Galaxy ecosystem.

## Background

Microbial communities (microbiomes) play fundamental roles across environmental, clinical, and engineered systems, influencing ecosystem functioning, host health, and biogeochemical cycles (Turnbaugh et al. 2007; Falkowski et al. 2008; Thompson et al. 2017). Despite their importance, a large fraction of microbial diversity remains poorly characterized, largely due to the historical reliance on culture-dependent microbiology. Traditional cultivation approaches capture only a small subset of microorganisms, as many taxa are recalcitrant to laboratory growth conditions, thereby limiting our understanding of community composition and functional potential (Epstein 2013; Lagier et al. 2018; Stewart 2012). The advent of high-throughput whole-genome sequencing (WGS) technologies has transformed microbiome research by enabling culture-independent investigation of microbial communities. In particular, whole metagenome shotgun sequencing (WMGS) has become a widely adopted approach to directly sequence the collective genomes present in a sample, providing simultaneous access to taxonomic diversity and functional gene content without prior cultivation (Quince et al. 2017). Best-practice guidelines and methodological frameworks have further standardized microbiome analyses and interpretation (Knight et al. 2018; Sharpton 2014). This shift has led to a rapid expansion in the volume and diversity of metagenomic datasets across a wide range of ecosystems, supported by large-scale genome catalogs (Nayfach, Roux, et al. 2021; Gurbich et al. 2023).

To extract biologically meaningful information from these complex datasets, genome-resolved metagenomics has emerged as a key analytical framework (Parks et al. 2017; Ayling et al. 2020; Hug et al. 2016). This approach enables the reconstruction of individual microbial genomes directly from metagenomic data, generating *metagenome-assembled genomes* (MAG), that represent draft genomes of uncultivated organisms. MAGs provide a powerful means to link phylogenetic identity with metabolic potential, offering new insights into the ecology, evolution, and functional roles of previously uncharacterized microorganisms (Ayling et al. 2020; Anantharaman et al. 2016).

Despite the critical role of MAGs in microbial research, a significant issue persists: many published studies do not provide publicly accessible or reproducible analysis pipelines. This lack of transparency not only undermines the reproducibility of results but also forces researchers to repeatedly develop custom pipelines, a time-consuming and error-prone process. Without standardized, shareable workflows, the scientific community faces redundant efforts, limited comparability between studies, and barriers to collaboration.

MAG reconstruction typically follows a series of core steps, including quality control of raw sequencing reads, assembly into contigs, binning of contigs into putative genomes, assessment of bin completeness and contamination, taxonomic classification, and functional annotation (Bowers et al. 2017). These steps form the foundational logic shared across most genome-resolved metagenomic pipelines, although their implementation and performance can vary depending on the tools and strategies used (Sczyrba et al. 2017; Meyer et al. 2022).

The choice of assembly and binning strategies, in particular, significantly impacts the quality of recovered MAGs. For example, reads can be assembled individually for each sample, grouped across biologically similar samples, or co-assembled across multiple samples. While co-assembly and grouped assembly can enhance the recovery of low-abundance genomes and improve overall completeness, they may also introduce artifacts such as cross-sample chimeras and increased contamination, especially in genetically heterogeneous samples (Haryono et al. 2022). In contrast, individual assemblies tend to produce MAGs with lower contamination and higher strain-level resolution, though they may miss genomes present at low abundance or only across some samples (Hofmeyr et al. 2020). For binning, co-binning (also referred to as multi-sample binning) is often considered optimal for MAG reconstruction, as it leverages coverage information across multiple samples to improve binning accuracy and genome completeness (Churcheward et al. 2022; H. Han et al. 2025; Kim et al. 2026). However, this approach is computationally intensive, requiring an all-vs-all read mapping step to generate abundance profiles across all samples. These methodological trade-offs, combined with the rapid evolution of tools, create a fragmented landscape of MAG workflows (Table 1). While the field benefits from continuous advancements in assemblers, binning algorithms, and refinement tools, this innovation introduces challenges for reproducibility, adaptability, and long-term maintenance. Even when workflows are shared, technical barriers frequently limit their adoption. Many workflows (Table 1) require expertise in workflow languages (e.g., Snakemake, Nextflow), or scripting (e.g., Bash, Python) as well as manual deployment on high-performance computing (HPC) clusters, introducing additional technical barriers and increasing the risk of configuration errors. Rigid, end-to-end designs, where any modification requires restarting the entire workflow, further complicate experimentation, while the lack of stepwise execution hinders troubleshooting and scalability.

**Table 1.** Non-exhaustive comparison of FAIRyMAGs with representative state-of-the-art metagenome-assembled genome (MAG) reconstruction workflows. The table summarizes key characteristics of selected MAG reconstruction pipelines, including supported target organisms, input data types, analytical steps (e.g., read quality control, assembly, binning, bin refinement, MAG quality assessment, taxonomic classification, and genome annotation), supported assembly strategies (e.g., individual, grouped, co-, and mixed assembly), and the underlying workflow management system. In addition, it compares workflow accessibility (e.g., web or command-line interfaces, modularity, and end-to-end execution), software availability (e.g., public repositories, workflow registries, containers, or package managers), and the availability of documentation, tutorials, and user support. An extended table with supported tools is available in the Supplementary Table 1.

| Platform / Workflow | Target Organisms | Input Type | Supported Analyses | Supported Approaches | Workflow Manager | Accessibility | Availability | Support & Documentation |
| --- | --- | --- | --- | --- | --- | --- | --- | --- |
| FAIRyMAGs | Bacteria | Short read | Read QC, Assembly, Contamination, Binning, Bin Refinement, MAGs Sample Pooling, MAGs QC, Taxonomy Classification, Genome Annotation | Individual, Co-, Grouped-, Mixed Assembly | Galaxy | Web-interface, Simple end-user modification, Independent runnable modules/sub-workflows, End-to-end runnable workflow | Public git repository, Workflow registry, Public server | Tutorial, User Support |
| Anvio's (Eren et al. 2021) | Bacteria, Eukaryotes, Viruses | Short read, Nanopore long read, Short & long read | Assembly, Binning, Bin Refinement, MAGs Sample Pooling, MAGs QC, Taxonomy Classification, Genome Annotation | Individual, Co-, Grouped-, Mixed Assembly<br>Automated assembly grouping, Multi-sample binning | Snakemake | Independent runnable modules/sub-workflows, Web-interface, Command-line interface, End-to-end runnable workflow | Public git repository | Tutorial, Documentation, User Support |
| ATLAS (Kieser et al. 2020) | Bacteria, Archaea | Short reads, long read (limited) | Read QC, Assembly, Binning, Bin Refinement, Taxonomy Classification, MAGs QC, MAGs Sample Pooling, Genome Annotation | Individual, Multi-sample binning. Mixed Assembly, Co-, | Snakemake | Independent runnable modules/sub-workflows, Command-line interface, End-to-end runnable workflow | Public git repository | Documentation, Tutorial, User Support |
| Aviary (Newell et al. 2024) | Bacteria, Archaea | Short read, Nanopore long read, Short & long read | Assembly, Binning, Bin Refinement, MAGs Sample Pooling, MAGs QC, Taxonomy Classification, Genome Annotation, Read QC | Individual, Co-assembly, Multi-sample binning, Automated assembly grouping | Snakemake | Independent runnable modules/sub-workflows, Command-line interface, End-to-end runnable workflow | Public git repository, Conda package | Documentation |
| MAGNETO (Churchward et al. 2022) | Bacteria, Archaea | Short read | Assembly, Binning, MAGs Sample Pooling, MAGs QC, Genome Annotation, Read QC, Taxonomy Classification, Bin Refinement | Individual, Co-assembly, Multi-sample binning, Automated assembly grouping | Snakemake | Command-line interface, End-to-end runnable workflow, Simple end-user modification | Public git repository, Workflow registry | Documentation |
| MetaflowX (Xia et al. 2025) | Bacteria, Eukaryotes, Viruses, Fungi | Hybrid, Short reads, assembled contigs | Read QC, Contamination, Assembly, Genome Annotation, Binning, Bin Refinement, MAGs Sample Pooling, Taxonomy Classification, MAGs QC | Individual, Co-, Multi-sample binning, automated assembly grouping, Hybrid assembly, Reference-base and reference free analyses | Nextflow | Command-line interface, End-to-end runnable workflow | Public git repository | Documentation |
| Metagenome-Assembled Genomes Orchestra (B et al. 2020) | Bacteria | Short read | Assembly, Binning, Taxonomy Classification, Genome Annotation | Individual assembly | Perl/bash scripts | Command-line interface, End-to-end runnable workflow | Container, Public git repository | Documentation |
| metagWGS (Mainguy et al. 2024) | Bacteria, Archaea | Short read, HiFi long read | Assembly, Binning, Bin Refinement, MAGs Sample Pooling, Taxonomy Classification, Genome Annotation | Individual, Co-, Grouped- assembly | Nextflow | Command-line interface, End-to-end runnable workflow | Public git repository | Documentation |
| METAWRAP (Uritskiy et al. 2018) | Bacteria, Archaea | Short read | Assembly, Binning, Bin Refinement, MAGs QC, Taxonomy Classification, Genome Annotation | Individual, Co-, Grouped- assembly, Multi-sample binning | Bash scripts | Independent runnable modules/sub-workflows, Command-line interface, End-to-end runnable workflow | Public git repository | Documentation |
| MGnify Genomes Generation pipeline (https://workflowhub.eu/workflows/884) | Bacteria, Archaea, Eukaryotes | Assembled short read | Read QC, Decontamination, Binning, Bin Refinement, Taxonomy Classification, MAG submission to ENA | Individual | Nextflow | Command-line interface, End-to-end runnable workflow | Public git repository, Workflow registry | Tutorial, Documentation, User Support |
| MuDoGer (Rocha, Coelho Kasmanas, et al. 2024) | Bacteria, Eukaryotes, Viruses | Short read | Assembly, Binning, MAGs Sample Pooling, MAGs QC, Taxonomy Classification, Genome Annotation | Individual | Bash scripts | Command-line interface, End-to-end runnable workflow | Public git repository | Documentation |
| nf-core/mag (Krakau et al. 2022) | Bacteria, Archaea | Short read, Nanopore long read, Short & long read | Assembly, Binning, Bin Refinement, Taxonomy Classification, Genome Annotation | Individual, Co-, Grouped- assembly, Multi-sample binning | Nextflow | Command-line interface, End-to-end runnable workflow | Public git repository, Workflow registry | Documentation, Tutorial, User Support |
| Metagenomics-Toolkit (Belmann et al. 2025) | Bacteria, Archaea | Short read, Nanopore long reads | Assembly, Binning, MAGs QC, Genome Annotation, Read QC, Contamination, Bin Refinement, Taxonomy Classification, Plasmid detection, Metabolic Modeling, Dereplication, Co-Occurrence, Fragment Recruitment | Individual, Multi-sample binning | Nextflow | Command-line interface, End-to-end runnable workflow | Public git repository | Documentation, Tutorial |

To address these challenges, we developed FAIRyMAGs, a FAIR-compliant, benchmarked and documented Galaxy pipeline for MAG reconstruction. Galaxy (The Galaxy Community 2026) is an open-source platform designed for FAIR data analysis. It features a user-friendly web interface, an automatable Application Programming Interface (API) and a powerful workflow management system, enabling users to combine bioinformatics tools into customizable workflows. Galaxy ensures full reproducibility by automatically capturing all parameters, inputs, and metadata required to repeat and validate analyses. Its intuitive workflow editor allows users to directly apply prebuilt workflows like FAIRyMAGs ones or adapt them to their specific needs. For example, to integrate new assemblers, binning algorithms, and downstream tools, as well as entire add-on workflows, such as viral MAG detection or antimicrobial resistance (AMR) detection pipelines. Indeed, because Galaxy tools and workflows are continuously developed and maintained by a large community, there is a constant influx of new tools and workflows. These can be used to extend and enhance the MAGs workflow.

A major advantage of Galaxy is its flexibility and scalability. Workflows can be executed on any Galaxy server, whether public or private, making it suitable for sensitive data requiring privacy. Major public Galaxy servers (e.g., usegalaxy.eu, usegalaxy.fr, usegalaxy.org.au, and usegalaxy.org) offer free access to substantial computing infrastructure, which is critical for resource-intensive tasks like metagenomic analysis. As the FAIRyMAGs workflows are openly available on two workflow registries (Dockstore (Yuen et al. 2021) and WorkflowHub (Gustafsson et al. 2025)), they can be directly imported and used on these servers or installed on any other Galaxy server with the required tools. With that, we eliminate the need for users to manage their own infrastructure, as the pipeline, including all required tools and reference data, is immediately accessible and ready to be used. To further reduce the use barrier, the FAIRyMAGs workflows are documented and supported by extensive training material freely available via the Galaxy Training Network (GTN) (Hiltemann et al. 2023). The material was already applied in three major metagenomic training events (<u>gxy.io/GTA2026-info</u>, <u>gxy.io/denbi-mg-2026</u>, <u>gxy.io/bari-mg-2026</u>) and received overall positive feedback.

To demonstrate the performance, robustness, and versatility of the FAIRyMAGs workflows in real-world applications, we reconstructed MAGs from four datasets representing distinct microbial ecosystems: an aeromicrobiome sampled from clouds and clear atmosphere, a termite head microbiome (unpublished), a macroalgal microbiome, and a bee gut microbiome. These datasets were selected to encompass a broad range of microbial communities, including both host-associated and environmental microbiomes, while covering varying levels of taxonomic complexity, sequencing depth, biomass availability, and expected genome diversity.

## Materials and methods

### FAIRyMAGs implementation

To reconstruct MAGs, FAIRyMAGs comprises a collection of 6 workflows, implemented in Galaxy (Fig. 1). Each workflow performs a distinct step required to generate MAGs and can be executed independently, enabling users to tailor their analysis according to their requirements. While those workflows constitute core functionalities required for MAGs building, they can also, by their generic design, be applied to various other genome analysis tasks. This approach promotes contributions from the broader Galaxy community.

**Figure 1.**
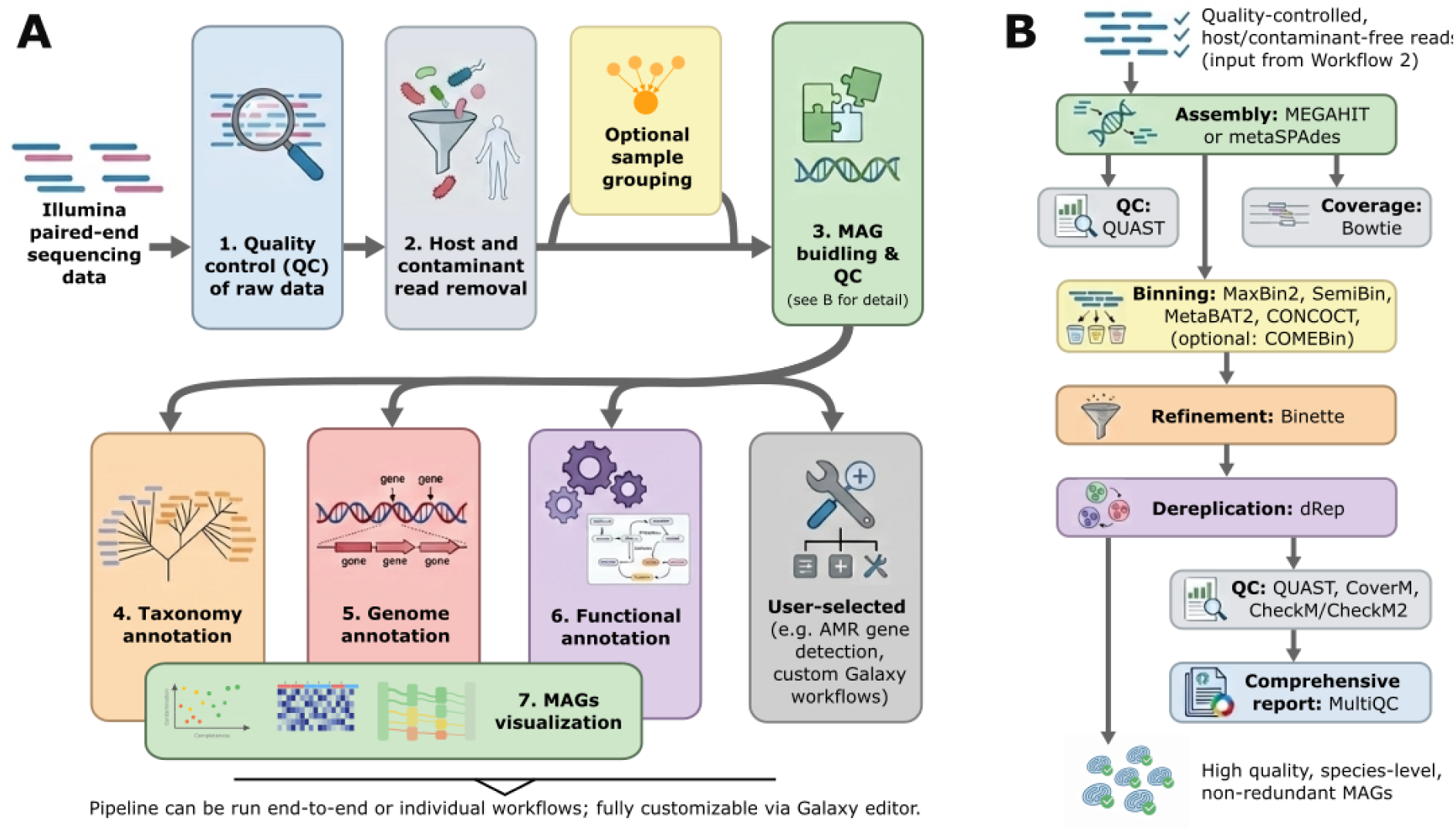
Interconnected workflows of the FAIRyMAGs pipeline for metagenome-assembled genome (MAG) construction and annotation. (A) Overview of the FAIRyMAGs pipeline. The FAIRyMAGs pipeline integrates multiple workflows to process Illumina paired-end sequencing data and generate high-quality MAGs. Workflow 1 performs quality control on raw sequencing data. Workflow 2 removes host and contaminant reads from the outputs of Workflow 1. Workflow 3 (detailed in panel B) is the core MAG building and quality control workflow, which can be executed directly on the outputs of Workflow 2 or after read grouping to enable grouped, co-assembly, or mixed assembly strategies. Following Workflow 3, parallel workflows are executed for downstream analyses: Workflow 4 for taxonomy annotation, Workflow 5 for genome annotation, Workflow 6 for functional annotation, and optional user-selected workflows available through the Intergalactic Workflow Commission (IWC), such as antimicrobial resistance (AMR) gene detection, or via custom Galaxy workflows. The pipeline can be run in its entirety or as individual workflows, with customizable parameters and adaptable workflows using the Galaxy workflow editor. (B) Detailed flowchart of Workflow 3: MAG building and quality control. This workflow begins from quality-controlled and host/contaminant-free reads. The first step involves metagenome assembly using MEGAHIT or metaSPAdes. The resulting contigs undergo quality assessment via QUAST and coverage evaluation by mapping original reads using Bowtie. Contigs are then binned using MaxBin2, SemiBin, MetaBAT2, CONCOCT, and optionally COMEBin tools, producing initial bins. These bins are refined using Binette. The refined bins are pooled together and processed with dRep for species-level dereplication to obtain non-redundant MAGs. The quality of these MAGs is assessed using CoverM, CheckM/CheckM2, and Quast. Finally, MultiQC aggregates the results into a comprehensive report.

The input data for FAIRyMAGs is paired-end sequencing data generated using Illumina. The datasets are preprocessed with Workflow 1, a workflow for quality control of paired-end short reads, followed by Workflow 2 for an host or contaminant read removal step. The host/contaminant read removal procedure in Workflow 2 can be repeated if multiple host genomes or potential contamination sources need to be filtered from the datasets. After preprocessing, the cleaned reads can be optionally grouped for co-assembly using the tools *fastq_groupmerge*. The grouped and cleaned reads are then processed in the main MAG building and quality control workflow (Workflow 3), which performs assembly, binning, refinement, dereplication, and quality control to generate good quality, species-level, non-redundant MAGs (Fig. 1B). The resulting MAGs are annotated via 3 parallel workflows: taxonomy (Workflow 4), genome (Workflow 5), and functional (Workflow 6). Optional workflows can also be used in addition to the main FAIRyMAGs workflows, such as the AMR gene detection workflow (Marin et al. 2025).

FAIRyMAGs provides a flexible and user-friendly framework for MAG reconstruction, enabling both comprehensive analyses and targeted investigations tailored to specific research needs. A critical aspect of robust annotation is the accuracy and currentness of reference data. To address this, FAIRyMAGs leverages Galaxy’s integrated Data Managers, which allow administrators to automatically download, install, and update essential reference databases. This ensures that users always work with complete, accurate, and up-to-date reference information. The workflows in FAIRyMAGs are configured to use well-maintained and reputable sources, such as GTDB (Parks et al. 2026) or AMRFinderPlus Reference Gene Catalog (Feldgarden et al. 2021), further enhancing the reliability of annotations. Additionally, Galaxy’s intuitive interface empowers users to select preferred reference databases or request the inclusion of specific datasets through their administrators, making the workflow highly adaptable to diverse research requirements.

#### Workflow 1: Quality control of raw data

The quality control (QC) workflow processes raw paired-end sequencing reads grouped into a collection to remove technical artifacts and low-quality bases before downstream analysis. Reads are trimmed and filtered using fastp (v1.1.0) (Chen et al. 2018), which performs adapter removal, base-quality trimming, and read length filtering. Quality metrics and preprocessing statistics are collected per sample and summarised across all datasets using MultiQC (v1.33) (Ewels et al. 2016) to provide an overview of read quality and filtering outcomes. The workflow outputs a collection of quality-controlled paired-end reads together with interactive QC reports generated by fastp and MultiQC. These filtered reads serve as input for the subsequent host and contaminant read removal workflow.

#### Workflow 2: Host and contaminant read removal

Workflow 2 reduces non-target sequence content to enrich microbial signals and reduce downstream assembly artifacts. This step is particularly important for host-associated microbiome datasets, as host-derived DNA may represent a substantial fraction of sequencing reads.

Quality-controlled reads from Workflow 1 are aligned against a host and/or additional optional contaminant reference genomes using Bowtie2 (v2.5.4) (Langmead and Salzberg 2012). The workflow can use locally installed reference genomes from the Galaxy server, benefiting from the large list of local reference genomes (160+ for usegalaxy.eu), or user-provided reference genomes. The read pairs mapping to the reference are filtered out, and only unmapped reads are retained for downstream analysis. The workflow also provides a summary of read recovery and removal across all samples using MultiQC (v1.33).

#### Optional read grouping for grouped-assembly

After preprocessing and before Workflow 3, the cleaned reads can optionally be merged into groups for grouped-assembly using the dedicated tool *fastq_groupmerge* (v1.0.2), which was developed as part of FAIRyMAGs. This tool allows flexible grouping of reads based on a user-defined sample sheet, enabling grouped assembly, co-assembly, or mixed assembly strategies in Workflow 3. Grouped assembly is typically recommended for biologically related samples, while co-assembly can help recover low-abundance MAGs (Pasolli et al. 2019). However, co-assembly should be used with caution, as pooling reads from multiple samples can introduce computational challenges during the assembly process. Because grouping is fully user-defined, the tool also supports more complex strategies, such as performing multiple alternative groupings or combining individual assemblies with selected grouped or co-assembly analyses.

#### Workflow 3: MAGs building and quality control

In Workflow 3, preprocessed and optionally grouped reads are assembled into contigs using either MEGAHIT (v1.2.9) (Li et al. 2015) or metaSPAdes (v4.2.0) (Nurk et al. 2017), MEGAHIT being the default choice.

To determine contig abundance (data needed for binning), reads are mapped to the contigs using Bowtie2 (v2.5.4). The resulting mapping file is sorted using Samtools *sort* (v2.0.7) (Danecek et al. 2021), and coverage depth is computed using MetaBAT2 (v2.17) (Kang et al. 2019) *calculate contig depths* tool.

Contigs are grouped into genome bins using four binning algorithms: MetaBAT2 (v2.17), MaxBin2 (v2.2.7) (Wu et al. 2016), CONCOCT (v1.1.0) (Alneberg et al. 2014), and SemiBin (v2.1.0) (Pan et al. 2022). In version 0.5 of FAIRyMAGs, COMEBin (v1.0.3) (Wang et al. 2024) was added as a fifth binner due to its strong performance in recent benchmarks (H. Han et al. 2025). Binning is performed using abundance information from the initially pooled samples. While multi-sample binning generally outperforms single-sample binning (Churcheward et al. 2022; H. Han et al. 2025), it also significantly increases runtime and memory usage. In particular, generating the cross-sample abundance profiles required by many multi-sample binning algorithms necessitates mapping the reads from each sample against contig assemblies representing all samples. Consequently, the computational cost of the mapping step increases rapidly with the number of samples, often becoming the primary bottleneck for large-scale studies. To balance efficiency and performance, a resource-efficient strategy is set as the default, though users can easily modify it to implement alternative approaches.

The resulting bins from the different binning tools are then refined and consolidated using Binette (v1.2.0) (Mainguy and Hoede 2024). After refinement, the MAGs are dereplicated across all samples using dRep (v3.6.2) (Olm et al. 2017), removing redundant genomes and selecting species-level representative MAGs based on quality metrics such as completeness and contamination. Since dRep internally supports only CheckM (v1.2.4) (Parks et al. 2015) for quality estimation, custom completeness and contamination metrics are provided instead, calculated using CheckM2 (v1.1.0) (Chklovski et al. 2023) and the CheckM2 *Diamond Reference Database v1.0.2* by default. For Binette and dRep, completeness and contamination thresholds are defined as workflow parameters, with default values of ≥75% completeness and ≤25% contamination, ensuring only high-quality MAGs are retained.

Finally, the quality of the dereplicated MAGs is estimated using CheckM2, with an optional CheckM *lineage_wf* step for additional validation. While CheckM has historically been the standard for MAG quality assessment using lineage-specific marker genes, CheckM2 uses a machine learning-based approach to improve predictions across diverse genomes. Assembly statistics are generated using QUAST (v5.3.0). The abundance of the MAGs in the samples is determined using CoverM (v0.7.0) (Aroney et al. 2025), by mapping a user-defined set of sample reads against the MAGs. This may include the individual samples used for the original assembly or any other sample set selected for abundance estimation. All quality metrics and sample abundance data are compiled and reported using MultiQC (v1.33), with results also provided in tabular format for downstream analysis.

#### Workflow 4: Taxonomy classification

In Workflow 4, the taxonomic classification of the reconstructed MAGs is performed using GTDB-Tk (v2.5.2) (Chaumeil et al. 2022) with the *GTDB database release 226* as default. GTDB-Tk assigns taxonomy based on the Genome Taxonomy Database (GTDB) (Parks et al. 2026), which provides a standardized, genome-based taxonomy and is widely recognized as the *de facto* standard for classifying bacterial and archaea MAGs (Parks et al. 2022).

However, the GTDB taxonomy differs from the NCBI taxonomy (Schoch et al. 2020), which remains widely used in downstream tools, databases, and comparative analyses. To bridge this gap and ensure compatibility, NCBI-GTDB-map (v0.1.9) (Youngblut et al. 2023) is used to map GTDB taxonomic identifiers to their corresponding NCBI Taxonomy identifiers. Following this, the Name2TaxID tool (v0.20.0) from the TaxonKit toolkit (Shen and Ren 2021) retrieves and generates the corresponding NCBI taxonomic names. This approach enables seamless integration of GTDB-based classifications with workflows, databases, and analyses that rely on the NCBI taxonomy, while preserving the original GTDB-based assignments.

Because GTDB-Tk is a computationally intensive tool requiring over 100 GB of memory per job, its application in high-throughput settings can result in extended processing times, particularly when executed through public Galaxy servers. Therefore, kMetaShot was additionally implemented as a rapid and lower-resource-demanding alternative for taxonomic classification of MAGs (Defazio et al. 2025). Both approaches can be applied in parallel, allowing comparison and validation of taxonomic assignments.

#### Workflow 5: Genome annotation

Workflow 5 performs structural annotation of MAGs, identifying gene locations, coding regions, and location of other structural elements. For comprehensive genome annotation, the workflow uses Bakta (v1.9.4) (Schwengers et al. 2021) with the *Bakta V5.1* and *AMRFinderPlus V3.12* databases as defaults. In addition to structural annotation, Bakta provides an initial layer of functional annotation by assigning curated gene names, product descriptions, EC numbers, and antimicrobial resistance genes. This annotation is subsequently extended in Workflow 6 with in-depth functional annotation.

For targeted annotation, the workflow employs ISEScan (v1.7.3) (Xie and Tang 2017) to identify insertion sequence and transposase, PlasmidFinder (v2.1.6) (Carattoli et al. 2014) with the *PlasmidFinder 2.1* database to detect plasmid replicons and plasmid-derived sequences, and Intregon Finder (v2.0.5) (Néron et al. 2022) to identify integrons and associated gene cassettes.

The workflow is based on an existing Galaxy workflow (Marin et al. 2025) extending summary tables for each tool, allowing standardized downstream analysis. These tables also serve as input for MultiQC, enabling unified visualization and comparison of all annotation results.

#### Workflow 6: Functional annotation

Workflow 6 extends the annotation from Workflow 5 (especially some functional annotation provided by Bakta by performing comprehensive functional annotation of MAGs using eggNOG (v2.1.13) (Cantalapiedra et al. 2021) with database version 5.0.2 and InterProScan (v5.59-91.0) (Jones et al. 2014) with database version 5.59-91.0 as default. The workflow adapted an existing workflow originally designed for protein sequences (Libouban and Bretaudeau 2025) to also process nucleotide sequences, significantly increasing its versatility. To further enhance functional insights, the workflow incorporates the kegg-pathways-completeness-tool (v1.3.0) (Richardson et al. 2023) which analyzes eggNOG results to calculate the pathway completeness of each MAG, providing a more comprehensive understanding of metabolic potential.

#### MAGs visualization

The final component of the FAIRyMAGs pipeline is the tool *MAGs-visualization*, an open-source Python-based command-line tool designed to transform complex MAG analysis outputs into publication-ready figures. It integrates and visualizes results from the Workflows 3-6, combining data from CheckM/Checkm2 for quality assessment, dRep for genome dereplication, GTDB for taxonomic classification, QUAST for assembly statistics, Bakta for genome annotation, CoverM for coverage analysis and relative abundance, and KEGG pathway annotations, with optional sample metadata inclusion.

*MAGs-visualization* produces core visualization, including completeness-contamination scatter plots for genome quality assessment, species-level clustering summaries from dRep results, taxonomic Sankey diagrams based on GTDB classifications, heatmaps of MAG abundance and detection patterns across samples, and functional annotation profiles reflecting KEGG pathway completeness. This tool enables to assess genome quality, explore taxonomic distributions, examine sample-specific MAG patterns, and compare metabolic potentials across MAGs and clusters, providing a comprehensive and interpretable summary of the entire FAIRyMAGs analysis pipeline.

### FAIR workflows

The FAIR principles (Findable, Accessible, Interoperable, and Reusable) provide a robust framework for enhancing the utility, reproducibility, and long-term usability of research objects, including data (Wilkinson et al. 2016), software (Chue Hong et al. 2022), or workflows (Visser et al. 2023). FAIRyMAGs was explicitly designed to adhere to these principles, fulfilling the 10 best practices for building FAIR workflows (Visser et al. 2023). To ensure portability and reproducibility, FAIRyMAGs leverages Galaxy as its workflow manager, providing a self-contained computational environment. The workflows employ standardized file formats (FASTA/FASTQ for sequence data, SAM/BAM for alignments, and GenBank/GFF3 for genomic annotations) ensuring seamless interoperability with other tools and platforms. The workflows are modular and include sensible default parameters, as described earlier.

All six workflows are publicly available in the GitHub repository of the Galaxy Intergalactic Workflow Commission (IWC), where they undergo rigorous review and testing with standardized datasets before each Galaxy release. Tools integrated into these workflows are semi-automatically updated to the latest Bioconda versions (Grüning et al. 2018), ensuring long-term compatibility. The workflows documented in this manuscript reflect the tool versions available at the time of submission. Each workflow follows best practices, including version control via GitHub releases and comprehensive metadata (e.g., licensing, authorship, and institutional affiliations). To maximize discoverability and accessibility, the workflows are automatically deposited in two major repositories (Dockstore (Yuen et al. 2021) and WorkflowHub (Gustafsson et al. 2025)) allowing users to install them on any Galaxy server with all needed tools. The workflows are also available and can be executed directly on four major Galaxy servers (usegalaxy.eu, usegalaxy.fr, usegalaxy.org.au, and usegalaxy.org), enabling federated use across distributed compute infrastructures.

A comprehensive tutorial for the full FAIRyMAGs pipeline is freely available through the Galaxy Training Network (GTN) (Hiltemann et al. 2023), featuring step-by-step guides, FAQs, test datasets, instructional videos, and interactive question boxes. Each individual workflow also has its own dedicated tutorial, all interconnected via a structured learning pathway (https://gxy.io/GTN:P00035).

Finally, to ensure full reproducibility and traceability, every workflow execution can generate a Research Object Crate (RO-Crate) (Soiland-Reyes et al. 2022), which archives all output data, parameters, tool versions, and workflow metadata in a machine-readable format. This feature enables complete documentation of computational analyses, supporting transparent and verifiable research.

### Use cases

To demonstrate the performance and versatility of the FAIRyMAGs workflows in real-world applications, we reconstructed MAGs from four datasets representing distinct microbial ecosystems: (i) aeromicrobiome sampled from clouds and clear atmosphere (BioProject: PRJEB54740), (ii) termite head microbiome (unpublished), (iii) macroalgal microbiome (BioProject: PRJNA915238), (iv) bee gut microbiome (BioProject: PRJNA977416). These datasets were selected to encompass a broad range of microbial communities, spanning both host-associated and environmental microbiomes with varying levels of community complexity, sequencing depth, and biomass availability. All datasets were processed using the complete FAIRyMAGs pipeline, comprising all six workflows (Workflow 1 v0.3, Workflow 2 v0.2, Workflow 3 v0.4, Workflow 4 v0.2, Workflow 5 v0.1, and Workflow 6 v0.3). Default parameters were used throughout, with dataset-specific adjustments applied only when required to accommodate the characteristics of individual datasets. These adjustments are described for each use case in the following section. MAGs results, including QUAST assembly statistics, dRep clustering output, CheckM2 quality assessment, GTDB taxonomy classification and BAKTA genome annotation, were further explored and visualized using Python (v3.14.9) within Jupyter Notebooks (v1.1.1). Data manipulation was performed using *pandas* (v3.0.3) (The pandas development team 2026; McKinney 2010) and *numpy* (v2.5.1) (Harris et al. 2020). Visualizations were created using *matplotlib* (v3.11.0) (Hunter 2007) and *searborn* (v0.13.2) (Waskom 2021), while *scikit-learn* (v1.9.0) (Pedregosa et al. 2011) was used for advanced statistical analyses, such as Principal Component Analysis (PCA). Additionally, the use cases were visualized using the *MAGs-visualization* tool (v0.0.11).

#### Aeromicrobiome

The aeromicrobiome dataset originates from the first study combining non-targeted metagenomic and metatranscriptomic approaches to investigate the functioning dynamics of airborne microbial communities in clouds compared to clear atmospheric conditions (Péguilhan et al. 2025). Samples were collected at the summit of the Puy de Dôme in 2019 and 2020, encompassing nine cloud events and five clear-air events. Sampling was conducted for 2 to 6 consecutive hours during the daytime using two to four high-flow-rate impingers operating in parallel. Nucleic acid were preserved in a filtered, autoclaved nucleic acid preservation (NAP) buffer, with collection volumes monitored hourly and adjusted with autoclaved ultrapure water as needed. The collected samples were filtered through 0.22 µm mixed cellulose ester (MCE) filters and DNA was extracted using the NucleoMag® DNA Water Kit. Half of each lysate was processed, and RNA was removed by RNase A treatment before DNA elution in DNase-free H₂O. Total DNA yields ranged from 42.6 to 838.7 ng, corresponding to airborne concentrations of 0.03 to 0.73 ng DNA/m³. DNA extracts from replicate samples of the same event were pooled, and 30 µL aliquots were submitted to GenoScreen (Lille, France) for shotgun metagenomic sequencing on an Illumina HiSeq platform (2 × 150 bp paired-end reads) (BioProject: PRJEB54740).

The raw sequencing data were processed through the FAIRyMAGs pipeline: quality control (Workflow 1) was followed by contaminant removal (Workflow 2) using the human reference genome (hg38, NCBI RefSeq assembly: GCF_000001405.26). For workflows 3, metaSPAdes was selected over MEGAHIT, based on prior benchmarking indicating improved performance for low- and medium-complexity datasets with shallow sequencing depth (Zhang et al. 2023). Additionally, non-default completeness and contamination thresholds were set for Binette and dRep to ≥0% completeness and ≤100% contamination, ensuring that all MAGs, including low-quality ones, were retained for downstream analysis. Workflows 4-6 were executed with default parameters to perform taxonomic and functional annotation of the reconstructed metagenome-assembled genomes (MAGs).

#### Termite Head Microbiome

The termite dataset consisted of 28 metagenomic libraries derived from two wood-dwelling termite species (*Cryptotermes secundus* and *Cryptotermes domesticus*), collected from mangrove habitats in the Darwin region, Australia. For each species, two independent colonies were sampled, with seven individuals per colony (five workers, one king, and one queen), resulting in a total of 28 samples. All colonies were maintained under standardized laboratory conditions following established protocols (Waidele et al. 2017).

Heads and guts from each individual were processed separately for DNA extraction using a bead-beating and organic extraction protocol (Waidele et al. 2017). Prior to extraction, ZymoBIOMICS Spike-in Control II (Low Microbial Load), containing *Truepera radiovictrix*, *Imtechella halotolerans*, and *Allobacillus halotolerans*, was added to each sample to monitor recovery efficiency and potential processing biases. Sequencing libraries were prepared using NEBNext® reagents with unique dual-index primers and sequenced on an Illumina NovaSeq 6000 platform (2 × 150 bp paired-end) at the Life & Brain Center (University Hospital Bonn, Germany).

A total of 56 metagenomic libraries were generated (two per individual, corresponding to head and gut samples). For this study, we focused exclusively on the 28 head-derived data. The reads underwent quality control (Workflow 1), followed by host read removal (Workflow 2) using the genomes of *C. secundus* (Csec_1.0; NCBI GenBank assembly accession GCA_977948025.2) and *C. domesticus* (unpublished in-house assembly). To enhance genome recovery, particularly for low-biomass head-associated microbiome samples, host-depleted reads were grouped by host species, colony, and caste prior to assembly using the *fastq_groupmerge* tool (v1.0.2), resulting in eight co-assembled read sets. The co-assembled reads were subsequently processed through Workflow 3 with dataset-specific parameter adjustments. Specifically, MEGAHIT was run with a minimum output contig length of 500 bp, and reconstructed MAGs were retained using thresholds of ≥70% completeness and ≤10% contamination. Workflows 4–6 were executed with default parameters. Reconstructed MAGs were mapped back to individual sample reads to estimate genome-level abundances.

#### Macroalgal Epiphytic Microbiome

This use case was derived from a study that investigated the functional characterization of epiphytic microbiomes (epibiomes) associated with three macroalgal species: two red algae (*Sphaerococcus coronopifolius* and *Asparagopsis taxiformis*) and one brown alga (*Halopteris scoparia*) (Lavecchia et al. 2024). The study aimed to explore the hypothesis that halogenated compounds produced by these algae influence interaction between the algae and their associated microbiomes. Three composite samples were collected from the harbor bay of Lagosteiros (37° 1′ 9.678′′ N 7° 55′ 49.584′′ W), in southwest Portugal via snorkeling in October 2016. Each sample contains at least 3 distinct individuals of the same algae species. Algae tissue samples were rinsed and centrifugated to isolate the microbial community.

Sequencing was performed on an Illumina NextSeq 500 platform with paired-end (2×75 bp) layout (BioProject PRJNA915238). The raw sequencing reads were processed through the FAIRyMAGs pipeline to reconstruct and analyze MAGs using default parameters.

#### Bee Gut Microbiome

The bee gut microbiome dataset was derived from a study investigating the effects of *Nosema ceranae* (a prevalent microsporidian parasite), a probiotic bacterium (*Pediococcus acidilactici*), and a neonicotinoid insecticide (thiamethoxam) on the honey bee gut microbiota (Sbaghdi et al. 2024). Worker honey bees (*Apis mellifera*, Buckfast genotype) were collected in September 2021 from three colonies located at the same apiary in Clermont-Ferrand, France. After CO_2_ anesthesia, groups of 80 worker bees were placed in Pain-type cages and subjected to seven experimental conditions, each replicated across the three colonies (21 samples total): (1) untreated control (C), (2) treatment with *P. acidilactici* (P), (3) chronic exposure to the thiamethoxam (T), (4) infection with *N. ceranae* (N), (5) *N. ceranae* infection followed by *P. acidilactici* treatment (NP), (6) *N. ceranae* infection followed by thiamethoxam exposure (NT), and (7) combined thiamethoxam exposure and *P. acidilactici* treatment (TP). Sixteen days post-treatment initiation, six whole guts (excluding crops) per sample were dissected, pooled, and processed to enrich bacterial DNA while minimizing host and parasite eukaryotic DNA contamination. DNA was extracted using a phenol-chloroform method, and 200 ng of DNA per sample was fragmented to ∼500 bp using a Covaris E220 (Covaris®). Paired-end libraries were prepared using the Illumina TruSeq® DNA Nano library kit and sequenced on a NovaSeq 6000 (Illumina®) to generate 2 × 150 bp reads, resulting in 21 metagenomic libraries (BioProject PRJNA977416).

The raw sequencing reads were processed through the FAIRyMAGs pipeline: quality control (Workflow 1) was followed by host read removal (Workflow 2) using the *Apis mellifera* reference genome (apiMel3.1, NCBI ID: GCF_003254395.2). Workflows 3–6 were executed with default parameters to reconstruct and analyze metagenome-assembled genomes (MAGs). Since the original study used MetaPhlAn4 (Blanco-Míguez et al. 2023) for read-based taxonomic profiling, which employs a custom taxonomy derived from the NCBI taxonomy, the GTDB-to-NCBI taxonomic mapping performed in Workflow 4 was of particular interest, to enable direct comparison of the FAIRyMAGs taxonomic assignments with the taxonomic profiles reported in the original study.

## Results

### Use cases

To assess FAIRyMAGs workflows on real-world datasets, MAGs were reconstructed from four distinct microbial ecosystems: (i) aeromicrobiome sampled from clouds and clear atmosphere (14 samples), (ii) termite head microbiome (28 samples), (iii) macroalgal epiphytic microbiome (3 samples), and (iv) bee gut microbiome (21 samples). These datasets exhibit distinct patterns in MAGs reconstruction (Table 2). A total of 230, 21, 167, and 161 MAGs were recovered from for the air, termite head, macroalgal epiphytic, and bee gut microbiomes, respectively. These MAGs clustered into 215, 14, 132, and 38 species-level clusters, with varying numbers of MAGs per cluster (e.g., 1.07 ± 0.46 for air, 4.24 ± 4.61 for bee gut) and different levels of completeness and contamination. The MIMAG classification (Bowers et al. 2017) of these clusters revealed notable differences in quality across use cases. For instance, the macroalgal epiphytic microbiome yielded the highest proportion of high-quality (HQ) clusters (72 clusters with >90% completeness and <5% contamination), followed by the bee gut microbiome (20 HQ clusters). In contrast, the aeromicrobiome had only 1 HQ cluster, reflecting its lower sequencing depth and its low biomass. Middle-quality (MQ) clusters (≥50% completeness, <10% contamination) were prevalent in all use cases, with the macroalgal epiphytic microbiome showing the highest number (124 MQ clusters). Low-quality (LQ) clusters (<50% completeness, <10% contamination) were only identified in the aeromicrobiome (178 LQ clusters), as MAGs with completeness <75% and contamination <25% were not reported for the three other use cases. These results highlight the variability in MAG quality across ecosystems, influenced by factors such as sequencing depth, sample complexity, and the stringency of quality thresholds.

**Table 2:**
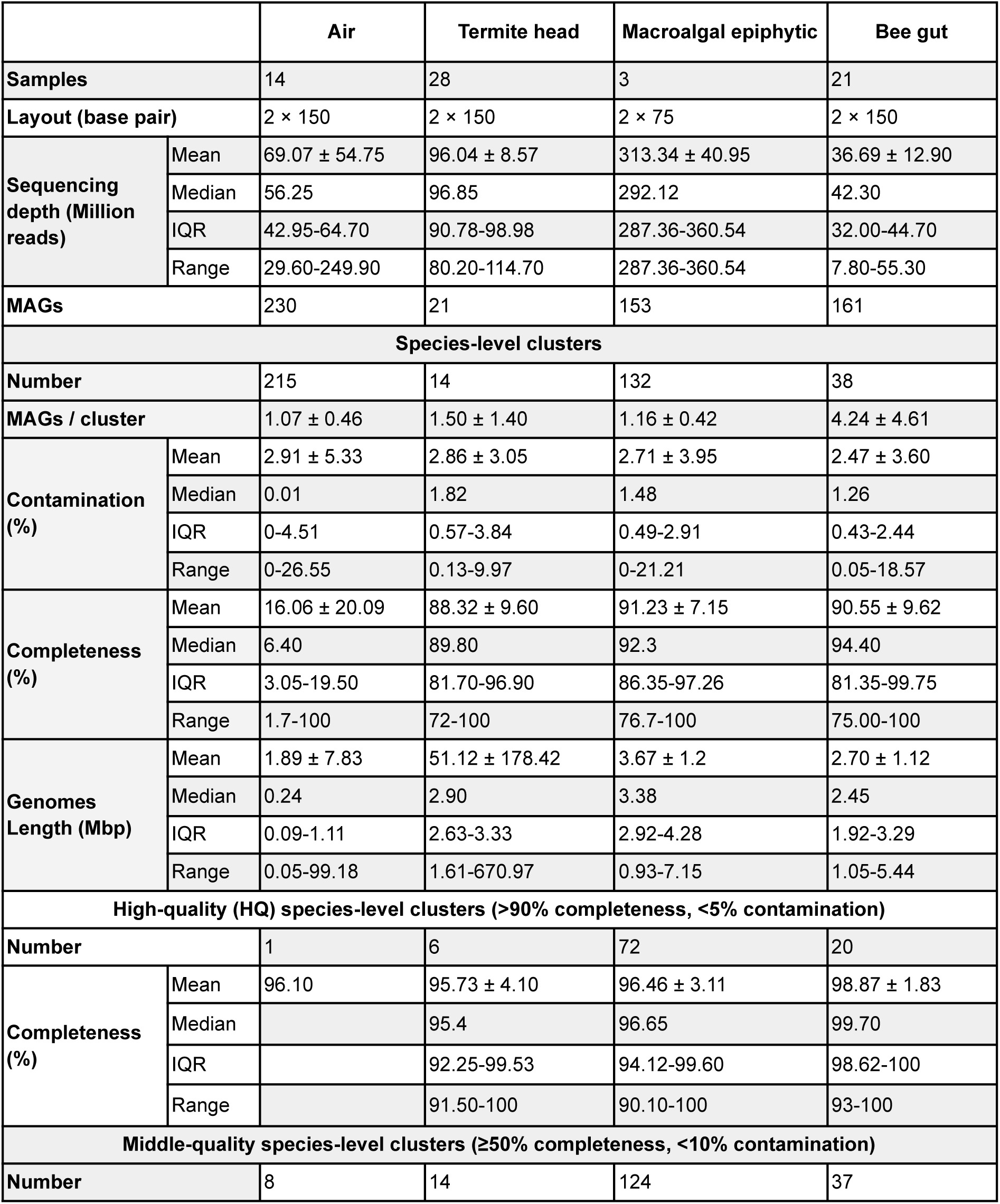
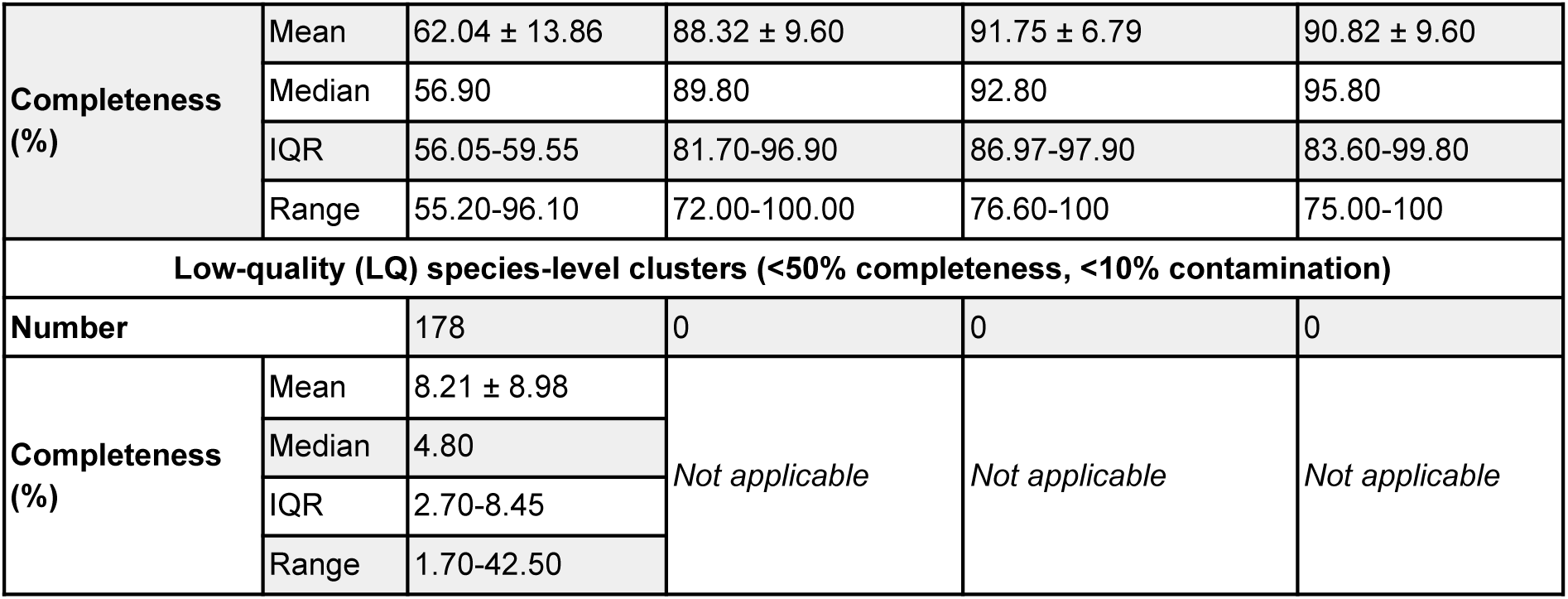
MAGs and species-level statistics for the four use cases (air, termite head, macroalgal epiphytic, and bee gut microbiomes). All statistics are computed for the representative genomes of the species-level clusters and reported as mean ± standard deviation, median, interquartile ranges (IQR), and range. High-quality (HQ), Middle-quality (MQ), and Low-quality (LQ) classifications follow the MIMAG standards (Bowers et al. 2017), where HQ clusters have >90% completeness and <5% contamination, MQ clusters have ≥50% completeness and <10% contamination, and LQ clusters have <50% completeness and <10% contamination. Clusters with contamination ≥10% are excluded from all quality groups, and HQ clusters are also included in the MQ clusters. *Note*: Statistics for all use cases except air are biased due to the completeness and contamination thresholds applied in Workflow 3, excluding MAGs with a completeness <75% or a contamination >25% (i.e. some MQ and all LQ clusters).

The taxonomic and functional annotations of the MQ species-level clusters further highlighted differences across the four use cases (Table 3). For instance, the bee gut microbiome exhibited the highest species-level classification (86.5% of clusters classified at the species level), while the aeromicrobiome had no species-level assignments. Genome and functional annotations, such as the number of KEGG modules, also varied, with the macroalgal epiphytic microbiome showing the highest mean number of modules (193.60 ± 20.75). The mapping rates to representative genomes of the MQ clusters ranged from 0.64 ± 0.40% for air to 67.82 ± 17.44% for termite head, indicating variations in the relative abundances of the recovered taxa. These differences in annotation completeness and mapping efficiency underscore the diversity in microbiome composition and assembly quality across ecosystems.

**Table 3:**
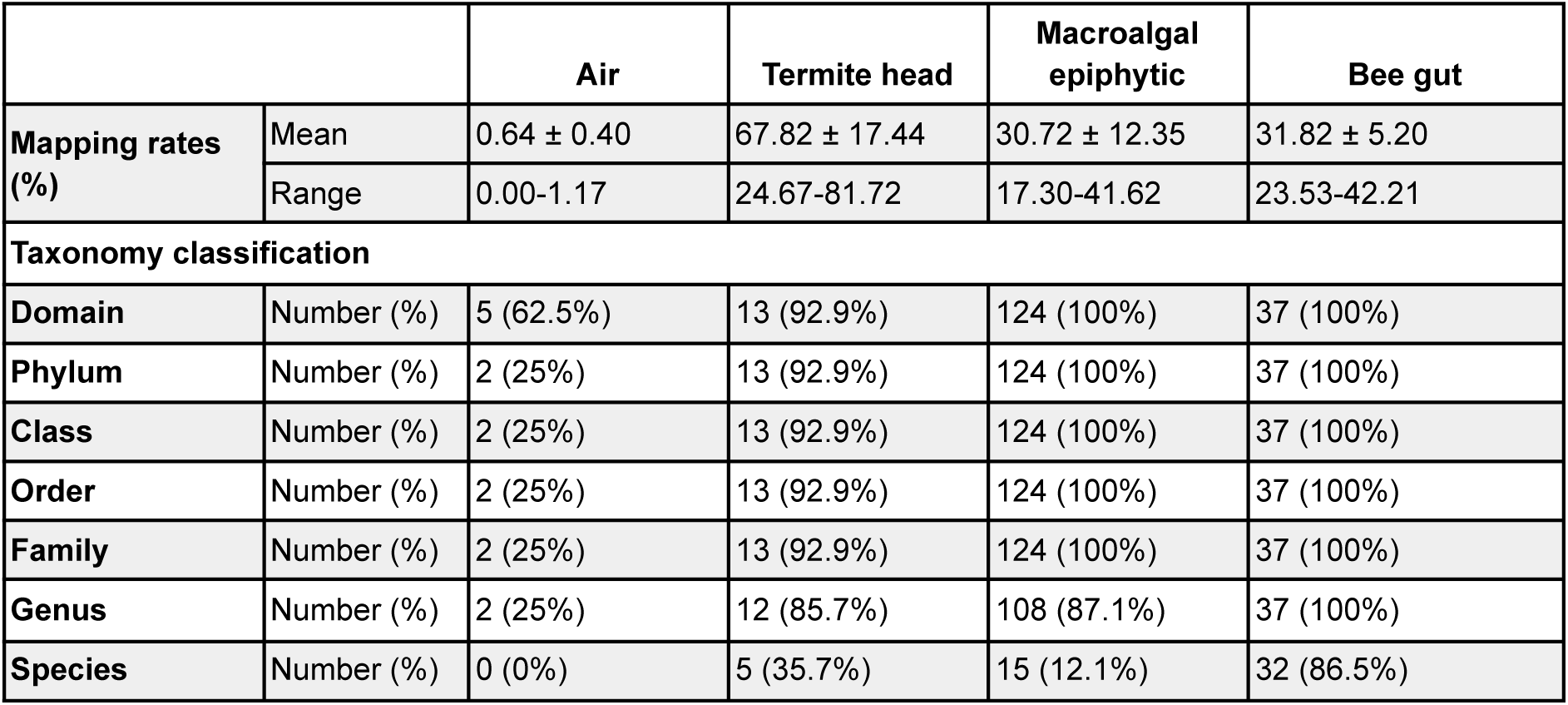
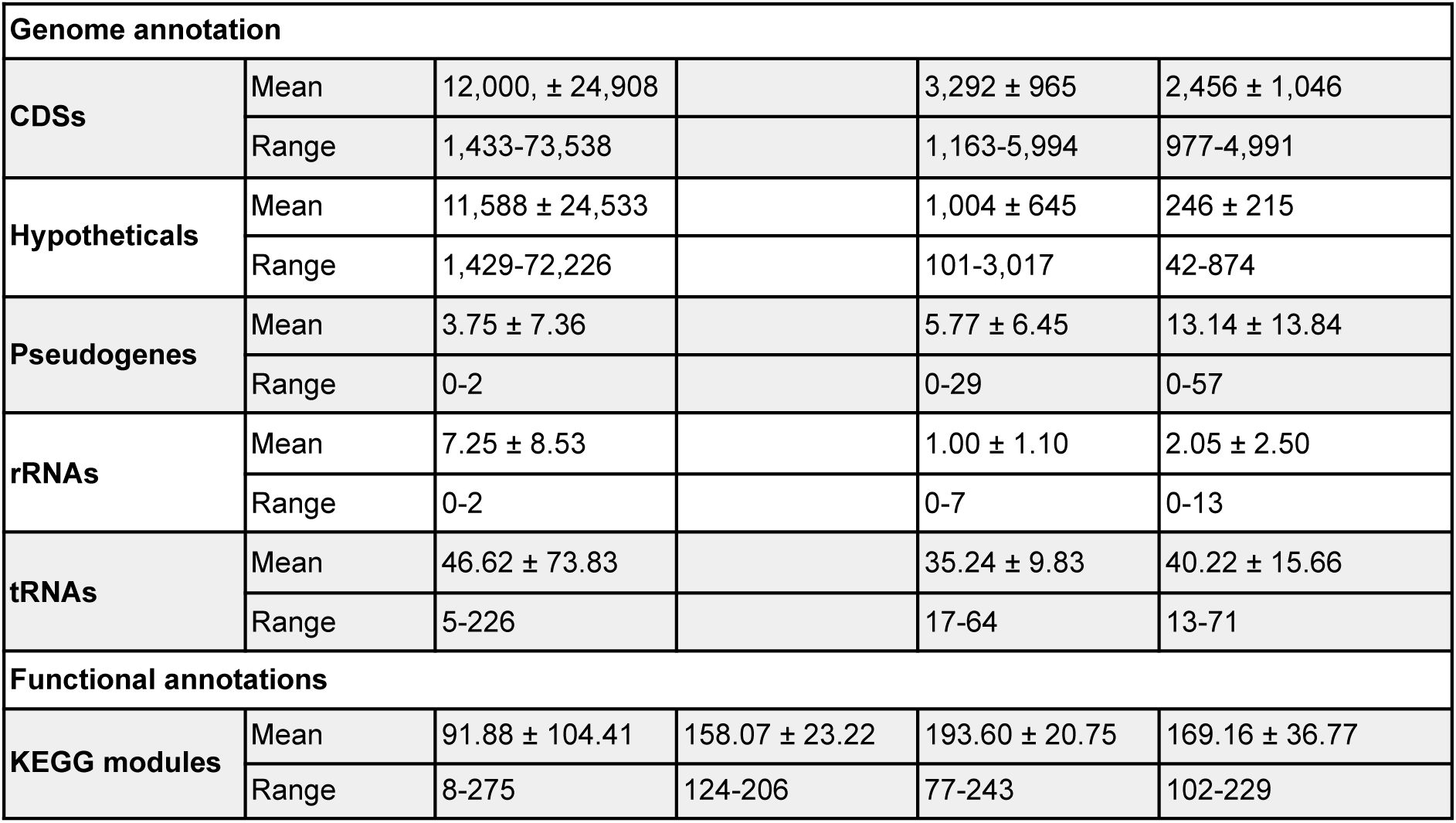
Statistics of taxonomic classification, genome annotations, and functional annotations of representative genomes of the middle-quality (MQ) species-level clusters for the four use cases. All statistics are reported as mean ± standard deviation and range. Mapping rates correspond to the percentage of reads (after Workflow 1 and 2) mapping to the representative genomes of the MQ species-level clusters. *Note*: Statistics for all use cases except air are biased due to the completeness (≥75%) and contamination thresholds applied in Workflow 3, excluding MAGs with a completeness <75%.

#### Aeromicrobiome

The aeromicrobiome dataset presented the greatest challenge, with 230 MAGs exhibiting a wide range of completeness (1.70-100%, median: 6.40%) and elevated contamination (up to 26.6%) (Table 2). Despite these challenges, 16 middle-quality (MQ) MAGs were retained across 8 species-level clusters, including 1 high-quality (HQ) MAGs (Supplementary Fig. 1, Table 2). Of these 8 MQ clusters, 5 representative MAGs are assigned to a domain (4 Bacteria and 1 Archaea), but only 2 could be taxonomically classified to the genus level (Table 3). Both belonged to the CAHJXG01 genus (Pseudomonadota phyla, Alphaproteobacteria class, Acetobacterales order, Acetobacteraceae family). When reads are mapped to these 8 representative MAGs (mapping rates: 0-1.17%), the CAHJXG01 genus accounts for 24.6 ± 32.67% of the relative abundance across samples. This high variability is explained by the absence of this genus in 1 clear atmosphere sample, its exclusive detection in 2 clear atmosphere samples (100% relative abundance), and its consistent presence in all cloud samples where it reached a relative abundance of 13.45 ± 5.86% (median: 12.48%, range: 4.00-24.65%). The representative MAGs (ERR9966625_bin_1350 and ERR9966620_bin_1469) exhibit strong structural differences despite similar contamination levels (3.94% and 3.73%, respectively), biassing the statistics in Table 2. ERR9966625_bin_1350 has a completeness of 96.1%, a genome length of 5.4 Mbp, and 5,189 CDSs (including 3,242 hypotheticals). In contrast, ERR9966620_bin_1469 had a completeness of 55.9%, a genome length of 43.5 Mbp (8.04 times larger), and 73,538 CDSs (14.17 times more), including 72,226 hypotheticals (22.28 times more). Additionally, ERR9966620_bin_1469 contains 21 rRNAs, 226 tRNAs, and 21 pseudogenes, compared to 0 rRNAs, 47 tRNAs, and 7 pseudogenes in ERR9966625_bin_1350. Both MAGs have similar KEGG module counts (230 and 275, respectively). These differences raise questions about taxonomic classification consistency. The ERR9966625_bin_1350 representative genome (reasonable genome size) was classified by GTDB-tk based on topology and ANI, while the ERR9966620_bin_1469 representative genome (large genome) was classified using relative evolutionary divergence (RED), with a good RED value (0.95053), indicating taxonomic novelty.

Among the middle-quality (MQ) species-level clusters, 6 clusters lacked complete taxonomic classification: 3 had no domain assignment, while 3 were classified only at the domain level (2 Bacteria, 1 Archaea) with unclassified phyla and lower taxonomic ranks. These clusters exhibited low completeness (55.20–59.70%), relatively high contamination (6.45–9.69%), and similar genome sizes (2.04 ± 0.58 Mbp; range: 1.30–2.74 Mbp). The Archaea cluster was detected in 7 of 14 samples, exclusively in cloud samples, while the remaining unclassified clusters were present in all but 3 samples (3 of the 5 clear atmosphere samples). Despite the lack of proper taxonomic classification, the representative genomes of these clusters were functionally annotated. They contained 2,878.50 ± 1,448.84 CDSs (range: 1,433–4,949), including high proportion of hypothetical proteins (2,872.00 ± 1,451.52 range: 1,429–4,945), and 16.67 ± 10.93 tRNAs (range: 5–37). KEGG module annotations averaged 38.33 ± 36.03 (range: 8–91). rRNAs were identified in 3 clusters: one bacterial cluster contained 16 rRNAs, while the two clusters without domain assignments had 8 and 13 rRNAs, respectively. These included 5S ribosomal RNAs (5–13 copies), 16S ribosomal RNAs (2 full, 5 partial, and 1 3’ truncated), and 23S ribosomal RNAs (1 full, 1 partial, and 1 3’ truncated). The identified rRNAs corresponded to RFAM families such as 5S ribosomal RNA (RF00001), Bacterial small subunit ribosomal RNA (RF00177), and Bacterial large subunit ribosomal RNA (RF02541), suggesting that the two clusters without domain assignments likely represent bacterial taxa, leaving one cluster without any domain assignment.

One notable low-quality (LQ) cluster has its representative genome identified as an unclassified Archaea with 100% completeness but 26.55% contamination, exceeding the threshold for higher quality classifications. The representative genome of this cluster had a genome size of 10.83 Mbp and was annotated with 20,836 CDSs, of which 20,823 were hypothetical proteins, indicating a highly uncharacterized genome. It contained 3 tRNAs but no rRNAs, and its functional potential was reflected in 169 KEGG modules. The high contamination level suggests potential chimerism or co-assembly of multiple strains, which may have impacted its classification and functional interpretation.

#### Termite head microbiome

The termite head microbiome initially yielded 33 MAGs grouped into 26 species-level clusters (Supplementary Fig. 2). Following quality filtering according to the MIMAG medium quality criteria (≥50% completeness and <10% contamination), the final dataset comprised 21 MAGs representing 14 MQ species-level clusters (Table 2). These genomes showed a mean completeness of 88.3 ± 9.6% and a mean contamination of 2.86 ± 3.05%. Six representative genomes fulfilled the high-quality criteria, with a mean completeness of 95.73% ± 4.10%, mean contamination of 1.18 ± 1.38%, and mean genome length of 4.35 ± 2.95 Mbp.

The recovered MAGs spanned five bacterial phyla: Spirochaetota, Bacteroidota, Deinococcota, Margulisbacteria, and Pseudomonadota (Supplementary Fig. 2). The representative genomes were assigned to members of the families Breznakiellaceae and Termititenacaceae, as well as the genus *Propionivibrio*, JAIRRR01 and JAITMY01. A genome assigned to the intracellular endosymbiont *Wolbachia sp000829315* was also recovered. Notably, the recovered MAGs also included genomes corresponding to the spike-in control organisms (*Truepera radiovictrix*), demonstrating its recovery across the analysed samples. Nine of the 14 representative genomes remained unclassified at the species level, including four of the six HQ representative genomes, highlighting the limited representation of termite-associated microorganisms in current reference databases. Based on KEGG annotation, 314 KEGG modules were identified across 14 MAGs, with a mean of 158.07± 23.22 modules per representative genome (median: 152.5; range: 124-206). Among the MAGs, PRPP biosynthesis (M00005), Pyruvate oxidation (M00307), β-oxidation, acyl-CoA synthesis (M00086), and Cysteine biosynthesis (M00021) were the most frequently detected KEGG models.

#### Macroalgal epiphytic microbiome

The macroalgal epiphytic microbiomes yielded 132 species-level clusters (Supplementary Fig. 3) and 124 representative MAGs from medium-quality (MQ) species-level clusters. These MAGs showed high diversity in completeness and contamination, with median values of 92.80% (IQR: 86.97–97.90%) and 1.30% (IQR: 0.39–2.54%), respectively (Table 2). Genome lengths ranged from 1.16 to 7.15 Mbp, with a median size of 3.38 Mbp (IQR: 2.92–4.15 Mbp).

Taxonomic classification of representative MAGs from the MQ species-level clusters revealed a broad taxonomic spectrum, with Pseudomonadota (67 MAGs), Bacteroidota (46 MAGs), Actinomycetota (7 MAGs), and Cyanobacteriota (5 MAGs) representing the most abundant phyla. At the genus level, the most frequently assigned genera were Aquimarina (7 MAGs), CALKXS01 (5 MAGs), Arenicella (4 MAGs), Marinagarivorans (4 MAGs), and Dokdonia (4 MAGs), while 17 MAGs remained unclassified at the genus level. At the species level, 120 representative MAGs remained unclassified, whereas the remaining MAGs were assigned to Cellulophaga lytica, Dokdonia sp947496725, Lacinutrix sp000211855, Maribacter litoralis, Marinagarivorans sp947494095, Polaribacter sp002005425, Pseudoalteromonas marina, Dokdonia sp000355805, Hellea sp020629875, Lentilitoribacter sp020628775, Maribacter_B vaceletii, Marinagarivorans algicola, Olleya sediminilitoris, Pseudoalteromonas atlantica, and Psychrobacter sp002377945.

Genome annotation identified an average of 3291.54 ± 965.32 CDSs per MAG (median: 3089.50; IQR: 2615.75–3815.00), including a median of 849.50 hypothetical proteins (IQR: 499.75–1356.25). CRISPR arrays ranged from 0 to 4 per MAG, with a mean of 0.19 ± 0.65. The number of annotated rRNA genes ranged from 0 to 7, with a mean of 1.00 ± 1.10, while tRNA counts showed a median of 34 (IQR: 28.00–41.00).

Based on the annotated CDSs, 375 KEGG modules were identified across the representative MAGs of the MQ species-level clusters. The most frequently detected modules included pyruvate oxidation (M00307), guanine ribonucleotide biosynthesis (M00050), tetrahydrofolate biosynthesis (M00126), and the dicarboxylate-hydroxybutyrate cycle (M00374). In contrast, methanogenesis (M00563), benzoyl-CoA degradation (M00541), the Bacillus anthracis pathogenicity signature (M00859), and glycosaminoglycan biosynthesis (M00057) were among the least frequently detected KEGG pathways.

#### Bee gut microbiome

The bee gut microbiome use case yielded 161 MAGs that clustered into 38 species-level clusters, with an average of 4.24 ± 4.61 MAGs per cluster (Table 2). The MIMAG classification revealed 20 HQ clusters and 37 MQ clusters, with no low-quality (LQ) clusters reported due to the exclusion of MAGs with contamination ≥10%.

The 159 reconstructed MAGs of bee-associated MQ species-level clusters exhibited high completeness (75–100%) and low contamination, with 54% of the MG clusters exceeding 90% completeness while remaining below 5% contamination (Fig. 3B). Taxonomy assignment identified 37 genera, with 148 MAGs (92.93%) further classified into 32 species (Table 2, Fig. 2, Fig. 3C). The microbiome is dominated by Pseudomonadota (75 MAGs across 22 species), Bacillota (58 MAGs across 9 species) and Bacillota_A (12 MAGs across 4 species) phyla, while Actinomycetota (8 MAGs) and Bacteroidota phyla (6 MAGs) are less abundant with only 1 species each (Fig. 3A and C). The five most prevalent genera are *Apilactobacillus* (19 MAGs, 2 species), *Frischella* (17 MAGs, 1 species), *Fructobacillus* (17

**Figure 2.**
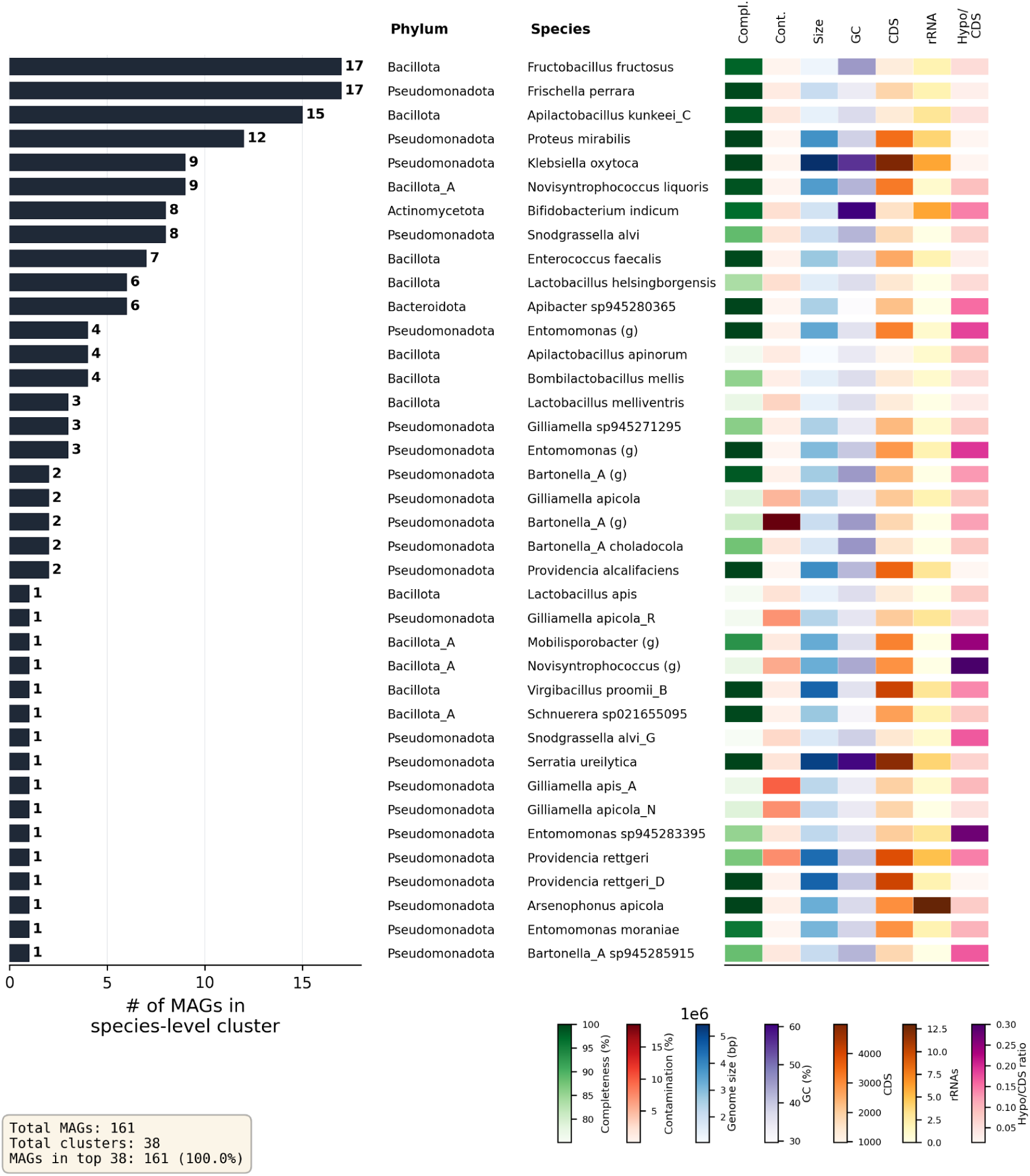
Distribution of species-level clusters in the bee gut microbiome use case. The horizontal bar plot shows species-level clusters, as defined by dRep secondary clustering (95% ANI), ranked by the number of MAGs per cluster. Bars represent cluster sizes, with the corresponding numbers of annotated MAGs. Taxonomic assignments (phylum and species) of the representative genomes are shown alongside each cluster. If no species-level assignment was possible, the next highest assigned taxonomic rank is displayed and indicated by its standard abbreviation (e.g., g. = genus). The heatmap panels summarize the quality and genomic characteristics of the representative genomes, including completeness (green), contamination (red), genome size (blue), GC content (purple), number of CDS (orange), number of rRNAs (orange), and the ratio of hypothetical proteins to CDS (pink). Summary statistics, including the total number of MAGs, the total number of species-level clusters, and the proportion of MAGs contained within the largest clusters, are shown in the lower-left corner of the figure.

**Figure 3.**
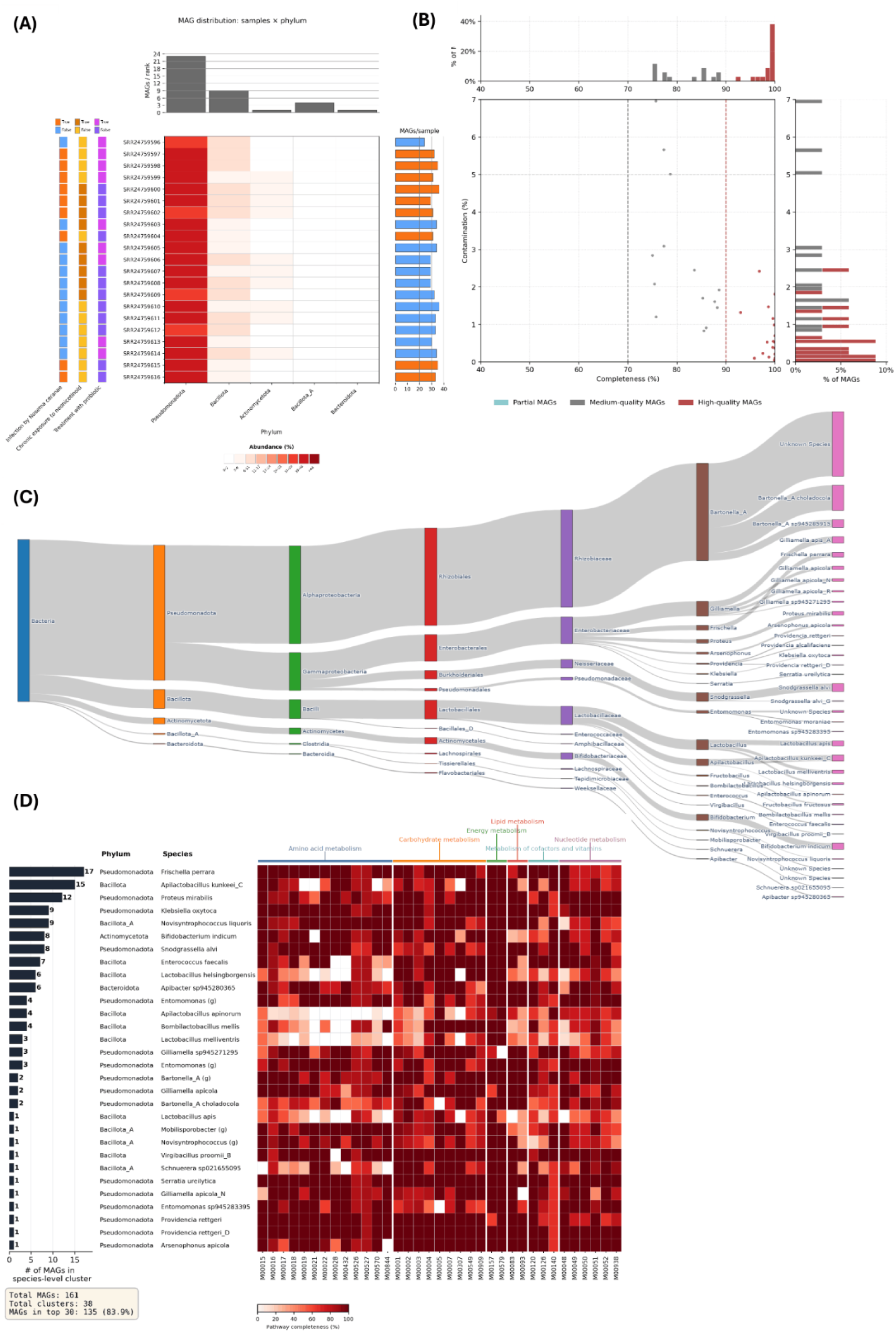
Recovered MAGs for the bee gut microbiome dataset. (A) Sample-by-phylum heatmap showing relative MAG abundance across samples, with associated metadata annotations indicating infection by *Nosema ceranae*, chronic exposure to neonicotinoids, and probiotic treatment. The bar chart on top shows the MAGs per taxonomic rank and the bar chart to the right shows the MAGs per sample with additional color coding based on infection by *Nosema ceranae.* (B) Completeness-contamination plot used for genome quality assessment, highlighting 70% and 90% completeness as well as 5 % contamination. Bar charts show the percentage of MAGs in this interval. (C) Sankey diagram illustrating abundance of MAGs at each taxonomic level, from domain to species (blue: domain, orange: phylum, green: class, red: order, purple: family, brown: genus, pink: species). (D) Heatmap of KEGG pathway completeness across representatives of the 30 largest species-level clusters, with pathways grouped by functional class. Modules were selected in “core” mode, which ranks all annotated KEGG modules by their mean completeness across all cluster representatives and displays the top-ranking ones (default: 35). This highlights pathways most broadly shared across the community.

MAGs, 1 species), *Proteus* (12 MAGs, 1 species), and *Lactobacillus* (10 MAGs, 3 species).

The relative abundance of representative MAGs of the MQ species-level clusters in the control group (C) revealed that non-core bacteria dominated the microbiota, with *Bartonella_A* species (2 over the 3 most abundant species) accounting for 38.2 ± 3.7% of the community. The core bee gut microbiota includes *Gilliamella apis* (8.3 ± 5.8%), *Snodgrassella alvi* (7.1 ± 2.8%), *Lactobacillus apis* (5.0 ± 1.0%), *Lactobacillus melliventris* (4.8 ± 3.0%), *Lactobacillus helsingborgensis* (3.4 ± 2.9%), *Gilliamella apicola* (2.6 ± 0.2%), *Gilliamella apicola_N* (2.7 ± 1.2%), and *Bombilactobacillus mellis* (0.5 ± 0.3%). Other notable species include *Bifidobacterium indicum* (7.0 ± 1.7%), *Frischella perrara* (3.7 ± 2.2%) and *Apilactobacillus kunkeei* (6.0 ± 1.8%). Across all experimental conditions, *Rhizobiaceae* (36.8 ± 5.8%)*, Enterobacteriaceae* (25.7 ± 7.8%) and *Lactobacillaceae* (18.5 ± 6.3%) are the most abundant families (Supplementary Fig. 4A). *Bartonella_A* (36.7 ± 5.8%) and *Gilliamella* (14.0 ± 4.7%) are the most abundant genera, with *Bartonella_A choladocola* (25.3 ± 5.1%) followed by *S. alvi* (8.0 ± 3.6%) as the most prevalent species. *Proteus mirabilis* shows a significant increase in infected honey bee gut (N, 5.3 ± 2.5%) compared to the control group (0.3 ± 0.02%). Some species, such as *Arsenophonus apicola* (21.4% in TP2), were detected only in specific samples, while remaining undetected or minimal in others (0.7 ± 0.4%). The PCA based on the relative abundance of species-level clusters (Supplementary Fig. 4B) reveals a condition-dependent structure, with samples from the same treatment group generally clustering together, though some variability was observed within certain groups (e.g., N1, N2, N3). Combined-stressor conditions (NT, NP, TP) occupied distinct positions relative to single-factor groups (C, T, P, N). Notably, infected samples (N, NT, NP) formed a separate cluster from non-infected samples, especially between N and C groups, indicating a clear distinction in community composition associated with *Nosema* infection. The species contribution magnitude (color-coded in the figure) further highlighted the influence of specific taxa, such as *Bartonella_A choladocola*, *Gilliamella apicola*, and *Snodgrassella alvi*, in driving these patterns. This pattern, alongside variation in phylum-level abundances (Fig. 3A), suggest microbiome compositional shifts potentially associated with y *Nosema ceranae* infection, chronic exposure to neonicotinoids, and probiotic treatment.

Across the representative MAGs of the 37 MQ species-level clusters, Bakta identified a mean of 2,456.2 ± 1,045.6 CDSs per genome (range 977-4,991), with hypothetical proteins accounting for a substantial fraction (245.1 ± 215.3; median 180.0) (Table 3). Non-coding features included means of 19.6 ncRNA regions, 15.6 ncRNAs, and 40.2 tRNA per genome.

Mobile/genomic-context signals were less abundant, with CRISPR arrays (mean 1.3; median 0; range 0–9), pseudogenes (mean 13.1; median 8; range 0–57), and rRNAs (mean 2.1; median 2.0; range 0–13). Replication/transfer origins (oriC, oriT, oriV) and tmRNAs were sparse, while gaps, oriV, and signal peptides were uniformly absent, suggesting either true absence or systematic under-detection. From the CDSs, 301 KEGG modules were reconstructed at varying completeness levels for 32 of the 38 representative MAGs. The number of modules decreased with increasing stringency: at 50% completeness, 31–171 modules per genome were identified (96.6 ± 37.3; median 94.5), dropping to 10–116 at 75% (58.8 ± 27.5; median 56.5) and 5–98 at 100% (40.3 ± 23.6; median 37.0). Notably, 90/301 modules had a completeness <50% in all 32 genomes (114 for <75%), while 6 modules were conserved across all 32 genomes at a completeness ≥50% (0 at 75%), including core pathways such as Guanine ribonucleotide biosynthesis (M00050), Nucleotide sugar biosynthesis (M00549), Glycolysis (M00001), C1-unit interconversion (M00140), Pentose phosphate pathway (M00004), and F-type ATPase (M00157). In these 32 species-level clusters (Fig. 3D), the core metabolic pathways, particularly those involved in amino acid and carbohydrate metabolism, are consistently present across clusters (Fig. 3D). In contrast, pathways related to lipid metabolism and cofactor/vitamin biosynthesis display greater variability, indicating differences in functional potential between clusters. This suggests that while central metabolic functions are largely conserved, certain taxa may exhibit more specialized metabolic capabilities.

## Discussion

### Workflow design

The developed workflow represents the first installment of a high-quality MAG generation pipeline within Galaxy. Despite the large number of MAG generation workflows proposed in recent years (Table 1), most pipelines converge on similar frameworks, largely due to the logical selection of best-performing tools for key steps such as assembly and binning. This convergence benefits the broader community by enabling continuous performance improvements at the level of individual tools.

The Galaxy-based implementation contributes to this development by offering several key advantages. These include access to large public computational resources, automatic tool updates, and the use of step-wise workflow execution by leveraging modular subworkflows for individual processing steps. In addition, Galaxy provides a range of features that enhance usability and reproducibility, including workflow versioning, detailed provenance tracking, workflow invocation graphs that allow users to follow individual execution steps, and seamless integration with data import and export tools.

Together, these aspects result in a best-practice pipeline with a low technical barrier, making it accessible to a wide range of users while remaining flexible for further optimization and expansion.

Technically, the workflow is comparable to those presented in Table 1, with the primary exception being the absence of co-binning in this initial version. Although co-binning can improve MAG recovery by leveraging information across multiple samples, it was omitted due to its high computational requirements. Performing cross-sample abundance profiling and comparisons across large metagenomic datasets can substantially increase both memory usage and runtime, potentially requiring orders of magnitude more resources than single-sample binning. To ensure accessibility and enable execution on publicly available computational infrastructures, we prioritized a workflow design that limits excessive memory and CPU requirements while maintaining broad applicability for users. This single-sample binning approach enables efficient processing of large numbers of independent samples, facilitating high-throughput MAG reconstruction studies without requiring extensive computational resources. However, a multi-sample binning version of the workflow is currently under development, targeting studies with a limited number of samples depending on the available computational resources.

Most publicly available workflows for generating MAGs provide an end-to-end analysis pipeline that starts from raw sequencing reads and, if successful, produces a set of reconstructed MAGs (compare Table 1). While these workflows simplify the analysis process, they typically operate as monolithic pipelines in which parameters can only be modified after the full workflow has completed. As a result, adjusting parameters often requires rerunning the entire pipeline, which can be computationally expensive and time-consuming.

However, MAG generation consists of multiple intermediate steps that may require user input or optimization at specific stages. For example, the quality control of raw sequencing reads often requires careful tuning of trimming parameters to achieve an optimal balance between removing low-quality bases and retaining as much usable data as possible. Overly strict trimming may remove informative reads, while overly permissive settings may allow low-quality sequences to propagate into downstream analyses. If such parameters are suboptimal, the effects may only become apparent at later stages of the workflow, such as assembly or binning, potentially reducing assembly quality or limiting the recovery of high-quality MAGs. In end-to-end pipelines, correcting these issues requires restarting the entire workflow with updated parameters. This uses unnecessary compute resources and hinders prototyping and parameter optimization on the user side.

Therefore, the FAIRyMAGs workflows were designed as modular components, each focusing on specific analytical tasks. These workflows can be executed independently, enabling parameter optimization and facilitating troubleshooting. The Galaxy workflow editor further supports flexible adaptation, allowing users to modify workflows by selecting different tool versions or replacing tools with alternatives from the Galaxy tool ecosystem. A wide range of such tools is maintained by the microGalaxy community (Nasr et al. 2024).

In this context, the FAIRyMAGs workflows serve as foundation for further development within the Galaxy community. Through updates distributed via the IWC, improvements can be propagated back to the workflows, ultimately benefiting the broader community.

Furthermore, the FAIRyMAGs workflows are currently limited to the reconstruction of archaeal and bacterial MAGs. This design reflects the focus of most existing binning algorithms, as well as the maturity of downstream quality assessment tools, such as CheckM2, and taxonomic classification frameworks, such as GTDB-Tk. Consequently, archaeal and bacterial genomes remain the primary targets of most current MAG reconstruction workflows (Table 1).

However, the Galaxy tool ecosystem provides a strong foundation for extending the FAIRyMAGs framework with dedicated workflows for the detection and analysis of viral and eukaryotic genomes. A viral genome reconstruction workflow is currently under development and will be available as an optional FAIRyMAGs module, integrating established Galaxy tools such as geNomad (Camargo et al. 2024) for virus identification, CheckV (Nayfach, Camargo, et al. 2021) for viral genome quality assessment, and Pharokka (Bouras et al. 2023) for genome annotation. Similarly, Galaxy already provides a broad range of tools for eukaryotic genome analysis, including Tiara (Karlicki et al. 2022) for sequence classification, MetaEuk (Levy Karin et al. 2020) for gene prediction, and BUSCO (Simão et al. 2015) for genome completeness assessment. These existing resources provide a solid foundation for future community-driven development and integration of dedicated eukaryotic MAG reconstruction workflows within the FAIRyMAGs ecosystem.

Another limitation is the use of short-read sequencing data as input for MAG reconstruction. While Workflow 3 was updated in version 5 to allow the inclusion of long reads as additional input for hybrid assembly with metaSPAdes, this functionality has not yet been thoroughly benchmarked. However, as hybrid assembly is also employed by several other workflows listed in Table 1, this approach is expected to be generally applicable.

Furthermore, several additional features could be incorporated into the FAIRyMAGs workflows, such as the option to manually refine MAGs using components from Anvi’o or to focus more on sample-wise analysis results instead of dereplicated study-wide MAG reporting. However, these features should be developed as independent downstream modules that extend the FAIRyMAGs ecosystem while preserving the modular design of the core workflows.

### Use cases

The use cases were independently processed by investigators with domain expertise in the respective microbiomes, some of them with limited knowledge of the Galaxy framework. All use case workflows could be executed successfully, from data upload to final MAGs and analysis results on public Galaxy servers. This demonstrates the technical usability of the workflows, their applicability across diverse experimental scenarios, and their potential for adaptation and further development by the community for MAG generation.

#### Aeromicrobiome

The aeromicrobiome dataset presented significant challenges in metagenome assembly, as evidenced by the low number of high-quality (HQ) and middle-quality (MQ) MAGs recovered. These difficulties are amplified by the low sequencing depth, a well-documented bias in MAGs reconstruction (Rocha et al. 2024; Treichel et al. 2026). Insufficient coverage likely hindered the assembly of HQ and MQ MAGs, suggesting greater sequencing efforts, similar to those employed in the macroalgal epiphytic use case, may be necessary in further studies to improve genome recovery. Additionally, co-assembly by condition (cloud vs clear atmosphere) could enhance MAG reconstruction, though this approach would require substantial computational resources, particularly in terms of RAM.

The LQ cluster with 100% completeness but 26.55% contamination raises important questions about completeness and contamination computations. While 100% completeness does not imply whole-genome recovery (Pellegrinetti et al. 2026), it does correlate with recovered functional signal, spanning all domains of metabolic functions (Eisenhofer et al. 2023). This suggests that even highly contaminated MAGs may retain valuable functional information, though their taxonomic and ecological interpretations should be treated with caution.

The taxonomic diversity observed in our MAGs was significantly lower than in the original study (Péguilhan et al. 2025), which used k-mer-based taxonomy classification on reads (Kraken2). The original study identified 32 distinct bacterial phyla, 159 orders, and 1,249 genera, with observed richness ranging from 532 to 826 genera per sample, alongside 8 eukaryotic phyla, 21 orders, and 54 genera, predominantly fungal. This discrepancy suggests that many taxa may have been missed in our MAGs due to low abundance, assembly limitations, or the presence of viral and small eukaryotic contributions, many of which may lack representation in public databases.

The CAHJXG01 genus identified in our MAGs appears to correspond to a genus within Rhodospirillales in the original study, though no direct equivalence exists between the two classifications. This discrepancy may stem from differences between NCBI and GTDB taxonomy databases. Regarding the unusually large representative genome (43.5 Mbp) of one of the two CAHJXG01 clusters, its annotation, particularly the high number of CDSs and hypothetical proteins, aligns with the known correlation between genome size and gene count (Kuo et al. 2009). However, typical CAHJXG01 genomes in GTDK are ∼5 Mbp, and bacterial genomes rarely exceed 14 Mbp (K. Han et al. 2013), suggesting that the 43.5 Mbp genome may represent an assembly artifact. However, the cluster contains 6 MAGs with genome sizes ranging from 8.97 to 100.69 Mbp (median: 59.04 Mbp), with all but one falling outside a reasonable range for bacteria. The representative genome was selected based on contamination levels (<5%) rather than size. This genome, despite its unusual characteristics, was taxonomically classified using relative evolutionary divergence, indicating potential taxonomic novelty, while the representative of the other CAHJXG01 cluster was classified by GTDB-tk based on topology and ANI. These differences highlight discrepancies in classification methods for genomes with atypical features.

For the unclassified clusters, 6 clusters lacked complete taxonomic classification: 3 were classified only at the domain level, 3 had no domain assignment. However, 2 of them could be confirmed as bacterial based on their rRNA content. The remaining cluster could potentially be eukaryotic, as GTDB annotations only include Archaea and Bacteria. To refine taxonomic assignments, eukaryotic taxonomic classification tools could be applied to this cluster. For the other clusters, more precise taxonomic assignments are needed. Potential approaches include using alternative tools like kMetaShot (Defazio et al. 2025) or leveraging rRNA-based classification for bacterial and archaeal clusters. Additionally, examining the reads mapped to each cluster and checking their taxonomic assignments via Kraken2 could provide further insights.

To better understand the reconstructed MAGs, large-scale k-mer searching could be conducting using Logan (Chikhi et al. 2025) to identify studies containing similar MAGs (e.g. (Drautz-Moses et al. 2022)) or sourmash branchwater (Irber et al. 2022) to identify matching genomes in public metagenomes, as done in (Lumian et al. 2024). These approaches could aid in taxonomic classification and help identify the potential origin and ecological roles of the organisms found in the atmosphere. However, improving MAG recovery for such understudied environments, characterized by low biomass, numerous viruses, small eukaryotes, and organisms with genomes not represented in public databases, remains a critical first step. Future efforts should focus on optimizing sequencing depth, assembly strategies, and taxonomic classification methods to enhance the recovery and interpretation of MAGs from complex, low-biomass environments.

#### Termite Head Microbiome

The termite head microbiome resulted in 14 representative MAGs after dereplication. Low biomass host associated microbiomes can bring challenges for metagenomic analyses because they are more susceptible to contamination and often contain limited microbial DNA relative to host associated reads (Eisenhofer et al. 2019). Nevertheless, all reconstructed MAGs exceeded the applied quality threshold (>70% completeness and <10% contamination), including six additionally fulfilled high-quality criteria (>90% completeness and <5% contamination), demonstrating that high genome quality was maintained despite the limited genome yield.

The recovered MAG collection included members of bacterial taxa previously associated with termite microbiomes, including *Spirochaetota*, *Bacteroidota*, *Margulisbacteria*, and *Pseudomonadota* (Arora et al. 2022; Salgado et al. 2024). Recovering of these lineages indicates that the reconstructed genomes represent biologically relevant members of the termite associated microbial community. In addition, successful reconstruction of the spike-in control organism *T. radiovictrix* provides independent validation that a genome of known origin present in the sequencing data could be accurately reconstructed throughout the workflow.

Several MAGs were assigned to environmental or taxonomically unresolved lineages. Their presence is not unexpected in host-associated metagenomic datasets and may represent microorganisms associated with the termite habitat or cuticle, low abundance community members, or the lineages that remained poorly represented in current reference databases. Overall, the termite use case shows that FAIRyMAGs can recover high-quality genome collections suitable for downstream genome-resolved analysis from low-biomass host associated metagenomic datasets.

#### Macroalgal Epiphytic Microbiome

The macroalgal epiphytic microbiome use case shows a reliable representation of this particular microbiome generated with the FAIRyMAGs workflows. Considering marine biodiversity and sequencing depth, and given that no widely recognized benchmark exists for the number of recoverable microbial genomes, the number of MAGs obtained here is quite low, notably lower than the 200 MAGs reported in the original paper. By contrast FAIRyMAGs was able to retrieve 72 and 124 representative MAGs of the high-quality and medium quality species-level clusters, respectively, representing the highest number among use cases and higher than the original paper that found 98 MQ MAGs. In contrast to FAIRyMAGs, the workflow used in the original publication did not include MAG dereplication. Therefore, the 98 MAGs identified in that study should also be considered potentially redundant. Nonetheless, the MAGs genomes length obtained is comparable with the expected prokaryotic genomes lengths (Rodríguez-Gijón et al. 2023). Overall, taxonomy profiling of MAGs is comparable with the original manuscript: works agree about the prevalence of Pseudomonadota (alias Proteobacteria) and Bacteroidota followed by Actinobacteriota, Mixococcota and Bdellovibriota. The same comparable taxonomic blueprint of MAGs can be reported at order rank with Flavobacteriales, Pseudomonadales, Enterobacteriales, Sphingomonadales.

#### Bee Gut Microbiome

The taxonomic profiling of the representative MAGs of our middle-quality species-level clusters revealed a high degree of concordance with the original study (Sbaghdi et al. 2024), after reconciling NCBI and GTDB taxonomies, 59.37% (19/32) of species matching those previously identified. All but three of these species (*Virgibacillus proomii_B*, *Apilactobacillus kunkeei_C*, *Frischella perrara*) are recognized as core members of the bee gut microbiome. However, 10 species reported in the original study are absent in our MAGs, including core taxa such as *Bifidobacterium asteroides*, *Bombilactobacillus mellifer*, *Lactobacillus kullabergensis*, and *Lactobacillus kimbladii*, as well as the dominant non-core species *Bartonella apis*. Potential matches for these missing taxa were identified in our dataset, such as *Bartonella_A* sp945285915 (putatively *Bartonella apis*), *Arsenophonus apicola* (putatively *Arsenophonus* spp.), *Providencia rettgeri_D* (putatively *Providencia* spp.), and *Serratia ureilytica* (putatively *Serratia marcescens*). Conversely, 19 species were uniquely found in our MAGs but not reported in the original study. This included *Enterococcus faecalis*, two unassigned genera in the *Lachnospiraceae* family, *Schnuerera* sp021655095, *Apibacter* sp945280365, four *Bartonella_A* spp., *Arsenophonus apicola*, four *Gilliamella* spp., two *Providencia* spp., *Serratia ureilytica*, and three *Entomomonas* spp. (two unassigned to species level). These discrepancies may reflect differences in taxonomic resolution between read-based (MetaPhlAn4) and MAG-based approaches, as well as variations in database coverage or annotation pipelines.

The most abundant families in our study (*Rhizobiaceae*, *Enterobacteriaceae*, *Lactobacillaceae*) align closely with those in the original study (*Bartonellaceae*, *Lactobacillaceae*), though the order of dominance differs slightly. Additionally, our analysis captured extra families not previously reported, potentially highlighting novel or underrepresented taxa in the bee gut microbiome.

The relative abundance patterns observed in our MAG-based analysis align with the original study in identifying dominant taxa, though with notable differences in their proportions. For instance, while *Bartonella apis* was the most abundant species in the original study (56.48 ± 6.1%), our analysis revealed *Bartonella_A* species (38.2 ± 3.7%) as the dominant group in the control samples, with *Bartonella_A choladocola* being the most prevalent. This discrepancy may stem from differences in taxonomic resolution (e.g., *Bartonellaceae* vs. *Rhizobiaceae*) and the inclusion of novel species not previously captured by read-based methods.

The core bee gut microbiota (e.g., *Gilliamella apis*, *Snodgrassella alvi*, *Lactobacillus spp.*) was consistently detected in both studies, though some core species (*Bifidobacterium asteroides*, *Bombilactobacillus mellifer*) were absent in our representative MAGs. This could reflect gaps in MAG recovery. Conversely, our analysis identified additional species (e.g., *Bifidobacterium indicum*, *Frischella perrara*, *Apilactobacillus kunkeei*) that were either minor or undetected in the original study, suggesting a broader detection range for MAG-based approaches.

The PCA results further highlight condition-dependent shifts in microbiome composition. In the original study, the infected group (N) separated clearly from the control (C), with *N. ceranae* and thiamethoxam (NT) or *P. acidilactici* (NP) treatments showing intermediate clustering. Our PCA (Supplementary Fig. 4B) similarly revealed a clear separation between C and N groups, with infected samples (N, NT, NP) forming a distinct cluster from non-infected samples, particularly between N and C. This consistency suggests that experimental conditions (e.g., *Nosema ceranae* infection, neonicotinoid exposure, probiotic treatment) drive reproducible shifts in microbiome structure. However, while the overall clustering patterns align, the specific taxa contributing to these patterns differ slightly between studies.

Overall, while abundance estimates and PCA trends are broadly comparable, the taxonomic identities and relative contributions of key species vary, likely due to methodological differences (MAGs vs. read-based taxonomy) and database dependencies. These findings emphasize the complementary nature of both approaches in capturing microbiome dynamics.

The functional annotation of the bee-associated representative MAGs reveals a highly conserved core metabolic capacity alongside specialized functional potential among species-level clusters. The presence of hypothetical proteins (245.1 ± 212.4 per genome) suggests a significant reservoir of uncharacterized genetic functions, which may include novel adaptations to the bee gut environment. The conservation of essential KEGG modules, such as glycolysis, pentose phosphate pathway, and nucleotide biosynthesis, across all 32 clusters aligns with the fundamental metabolic demands of microbial life, ensuring energy production, biosynthetic capabilities, and cellular maintenance. These pathways are critical for symbiotic stability in the bee gut, where microbes must efficiently utilize host-derived nutrients. The variability in lipid metabolism and cofactor/vitamin biosynthesis pathways points to functional differentiation among taxa, potentially reflecting niche partitioning or adaptive responses to environmental stressors (e.g., pesticide exposure or pathogen infection). For instance, the heterogeneity in KEGG module completeness (e.g., 90/301 modules with <50% completeness) may indicate taxon-specific metabolic specializations, such as the ability to degrade complex carbohydrates or synthesize vitamins that complement host nutrition. The absence of certain genomic features (e.g., oriV, signal peptides) could also suggest streamlined genomes in some taxa, a common trait in host-associated microbes that rely on the host for specific functions. Unlike the original study, which focused solely on taxonomic identification, our MAG-based approach provides critical functional insights, revealing the metabolic potential of the bee gut microbiome. These findings highlight the dual nature of the bee gut microbiome: a stable core that supports essential metabolic processes and a variable periphery that may confer resilience or adaptive advantages under changing conditions. This functional plasticity could be key to understanding how bee gut communities respond to anthropogenic stressors like pesticides or probiotic interventions, as seen in the PCA-based shifts in community composition.

## Conclusion

Similar to the goal of the Galaxy framework to make advanced computational analyses accessible to anyone, this work aimed to facilitate the generation of high-quality MAGs from both user-generated and public datasets without requiring programming expertise or dedicated HPC infrastructure. To achieve this, the FAIRyMAGs workflows provide the first comprehensive MAG generation framework in Galaxy that is comparable to established MAG workflows, while offering a flexible foundation for community-driven adaptation and further improvement.

The successful application of the workflows to diverse real-world use cases demonstrated their usability across different microbiome scenarios and their ability to generate MAGs comparable to those reported in published studies. We envision that FAIRyMAGs will continue to evolve through community contributions, enabling adaptation to specific research needs and integration of emerging tools within the rapidly expanding MAG generation landscape.

In light of recent developments that facilitate Galaxy tool development through LLM-supported tool wrapping, automated port of workflows from nf-core and Snakemake, as well as fully model-driven execution of tasks through the Galaxy MCP layer (https://galaxyproject.org/community/sig/ml-ai-across-galaxy), it is conceivable that MAG workflows currently developed independently by workflow-engine-specific communities will increasingly converge towards a unified framework. Such a framework could enable the consistent integration of the latest advances in MAG reconstruction while reducing fragmentation across different workflow ecosystems.

Building on the foundation established by the FAIRyMAGs workflows, Galaxy can play a vital role in supporting these unifying and expanding developments by providing the infrastructure to execute complex workflows using public computational resources, investigate intermediate and final results through integrated analysis tools and visualizations, and prototype new workflow logic, subworkflows, individual tools, and reference databases.

Ultimately, these efforts will support broader access to adaptable MAG generation workflows and contribute to improving our understanding of the biological dark matter represented by uncultivated microbial diversity.

## Declarations

### Availability of data and materials

The FAIRyMAGs workflows are published in IWC:

1. Workflow 1: Quality control of raw data

○ GitHub: https://github.com/galaxyproject/iwc/tree/main/workflows/read-preprocessing/short-read-qc-trimming
○ IWC: https://iwc.galaxyproject.org/workflow/short-read-qc-trimming-main/
○ DOI: 10.5281/zenodo.17805254
○ Workflowhub: https://workflowhub.eu/workflows/2025?version=4
○ Dockstore: https://dockstore.org/workflows/github.com/iwc-workflows/short-read-qc-trimming/main:main?tab=info
2. Workflow 2: Host and contaminant read removal

○ GitHub: https://github.com/galaxyproject/iwc/tree/main/workflows/microbiome/host-contamination-removal/host-contamination-removal-short-reads
○ IWC: https://iwc.galaxyproject.org/workflow/host-contamination-removal-short-reads-main/
○ DOI: 10.5281/zenodo.17832962
○ Workflowhub: https://workflowhub.eu/workflows/2026?version=4
○ Dockstore: https://dockstore.org/workflows/github.com/iwc-workflows/host-contamination-removal-short-reads/main:main?tab=info
3. Workflow 3: MAGs building and quality control

○ GitHub: https://github.com/galaxyproject/iwc/tree/main/workflows/microbiome/mags-building
○ IWC: https://iwc.galaxyproject.org/workflow/mags-building-main/
○ DOI: 10.5281/zenodo.15303665
○ Workflowhub: https://workflowhub.eu/workflows/1352?version=6
○ Dockstore: https://dockstore.org/workflows/github.com/iwc-workflows/mags-building/main:main?tab=info
4. Workflow 5: Taxonomy classification

○ GitHub: https://github.com/galaxyproject/iwc/tree/main/workflows/microbiome/mags-taxonomy-annotation
○ IWC: https://iwc.galaxyproject.org/workflow/mags-taxonomy-annotation-main/
○ DOI: 10.5281/zenodo.18853307
○ Workflowhub: https://workflowhub.eu/workflows/2099?version=3
○ Dockstore: https://dockstore.org/workflows/github.com/iwc-workflows/mags-taxonomy-annotation/main:main?tab=info
5. Workflow 4: Genome annotation

○ GitHub: https://github.com/galaxyproject/iwc/tree/main/workflows/microbiome/mag-genome-annotation-parallel
○ IWC: https://iwc.galaxyproject.org/workflow/mag-genome-annotation-parallel-main/
○ DOI: 10.5281/zenodo.19111963
○ Workflowhub: https://workflowhub.eu/workflows/2115?version=3
○ Dockstore: https://dockstore.org/workflows/github.com/iwc-workflows/mag-genome-annotation-parallel/main:main?tab=info
6. Workflow 6: Functional annotation
○ GitHub: https://github.com/galaxyproject/iwc/tree/main/workflows/genome_annotation/functional-annotation/functional-annotation-of-sequences
○ IWC: https://iwc.galaxyproject.org/workflow/functional-annotation-of-sequences-main/
○ DOI: 10.5281/zenodo.19187248
○ Workflowhub: https://workflowhub.eu/workflows/2118?version=2
○ Dockstore: https://dockstore.org/workflows/github.com/iwc-workflows/functional-annotation-of-sequences/main:main?tab=info

The tool MAGs-visualization is available via:

- GitHub: https://github.com/usegalaxy-eu/MAGs-visualization
- Conda: https://bioconda.github.io/recipes/mags-visualization/README.html
- Galaxy Wrapper for each plotting subcommand:

a. Comp-conta: https://usegalaxy.eu/?tool_id=toolshed.g2.bx.psu.edu/repos/iuc/mags_visualization_comp_conta/mags_visualization_comp_conta/0.0.11+galaxy0
b. Sample-heatmap: https://usegalaxy.eu/?tool_id=toolshed.g2.bx.psu.edu/repos/iuc/mags_visualization_sample_heatmap/mags_visualization_sample_heatmap/0.0.11+galaxy0
c. dRep-cluster-annot: https://usegalaxy.eu/?tool_id=toolshed.g2.bx.psu.edu/repos/iuc/mags_visualization_drep_cluster_annot/mags_visualization_drep_cluster_annot/0.0.11+galaxy0
d. dRep-cluster-func: https://usegalaxy.eu/?tool_id=toolshed.g2.bx.psu.edu/repos/iuc/mags_visualization_drep_cluster_func/mags_visualization_drep_cluster_func/0.0.11+galaxy0
e. Pathway-module-heatmap: https://usegalaxy.eu/?tool_id=toolshed.g2.bx.psu.edu/repos/iuc/mags_visualization_pathway_module_heatmap/mags_visualization_pathway_module_heat map/0.0.11+galaxy0
f. Taxa-sankey: https://usegalaxy.eu/?tool_id=toolshed.g2.bx.psu.edu/repos/iuc/mags_visualization_taxa_sankey/mags_visualization_taxa_sankey/0.0.11+galaxy0

The scripts to analyse the data as well as the results of the use cases are available in the GitHub repository: https://github.com/usegalaxy-eu/FAIRyMAGs

The raw sequencing reads for the use cases are publicly available through the Sequence Read Archive (SRA) under their respective BioProject IDs, except for the termite head microbiome dataset, which will be deposited upon publication of the associated manuscript currently in preparation:

1. Aeromicrobiome (BioProject: PRJEB54740)
2. Macroalgal microbiome (BioProject: PRJNA915238)
3. Bee gut microbiome (BioProject: PRJNA977416)

License: MIT

Any restrictions to use by non-academics: None

## Funding

German Federal Ministry of Education and Research, BMBF [031 A538A de.NBI-RBC]; Ministry of Science, Research and the Arts Baden-Württemberg (MWK) within the framework of LIBIS/de.NBI Freiburg.

de.NBI Cloud within the German Network for Bioinformatics Infrastructure (de.NBI) and ELIXIR-DE (Forschungszentrum Jülich and W-de.NBI-001, W-de.NBI-004, W-de.NBI-008, W-de.NBI-010, W-de.NBI-013, W-de.NBI-014, W-de.NBI-016, W-de.NBI-022).

Future Investment Program subsidized by the National Research Agency, ANR-11-INBS-0013 European Molecular Biology Laboratory core funds.

## Acknowledgements

The authors thank the expert reviewers who supported the analysis of the MAG use cases in addition to the authors of this manuscript: Nicolas Blot and Ivan Wawrzyniak (Laboratoire Microorganismes: Génome et Environnement (LMGE), France), and Antonio Placido (National Research Council, Italy).

The authors would like to thank Nils Kleinbölting and Peter Belmann (IBG-5: Computational Metagenomics, Institute of Bio- and Geosciences (IBG), Forschungszentrum Jülich GmbH, Germany), Hugo Lefeuvre and Samuel Chaffron (Nantes Université, École Centrale Nantes, CNRS, LS2N, UMR 6004, France), Diego Alvarez Saravia (Departamento de Ingeniería en Computación, Facultad de Ingeniería, Universidad de Magallanes, Chile), and James A. Fellows Yates (Microbiome Sciences Group, Department of Archaeogenetics, Max Planck Institute for Evolutionary Anthropology, Germany; Research Group Archaeogenetics, Leibniz Institute for Natural Product Research and Infection Biology Hans Knöll Institute, Germany) for their constructive feedback, valuable discussions, and helpful suggestions during the preparation of this manuscript. Their expertise and input contributed to improving the clarity and quality of the presented work.

Use of Generative AI. During the preparation of this manuscript, the authors used ChatGPT (OpenAI, GPT-5.5-mini) and Le Chat (Mistral, Mistral Medium 3.5) to support English language editing and refinement, including spelling corrections, grammar improvements, and suggestions for clearer scientific phrasing. In addition, Open WebUI (Freiburg AI infrastructure) with the OpenAI open-weight model gpt-oss-120b and Claude code with GPT-5.3-Codex model were used to assist with formatting, restructuring, and improving the readability of analysis scripts. These tools were not used to generate analysis workflows, develop computational methods, produce scientific results, or formulate scientific conclusions. All AI-assisted suggestions were reviewed, verified, and adapted by the authors where appropriate.

