## Supplementary Fig. 3 for "FAIRyMAGs - a series of FAIR Galaxy workflows for the generation of metagenome assembled genomes"

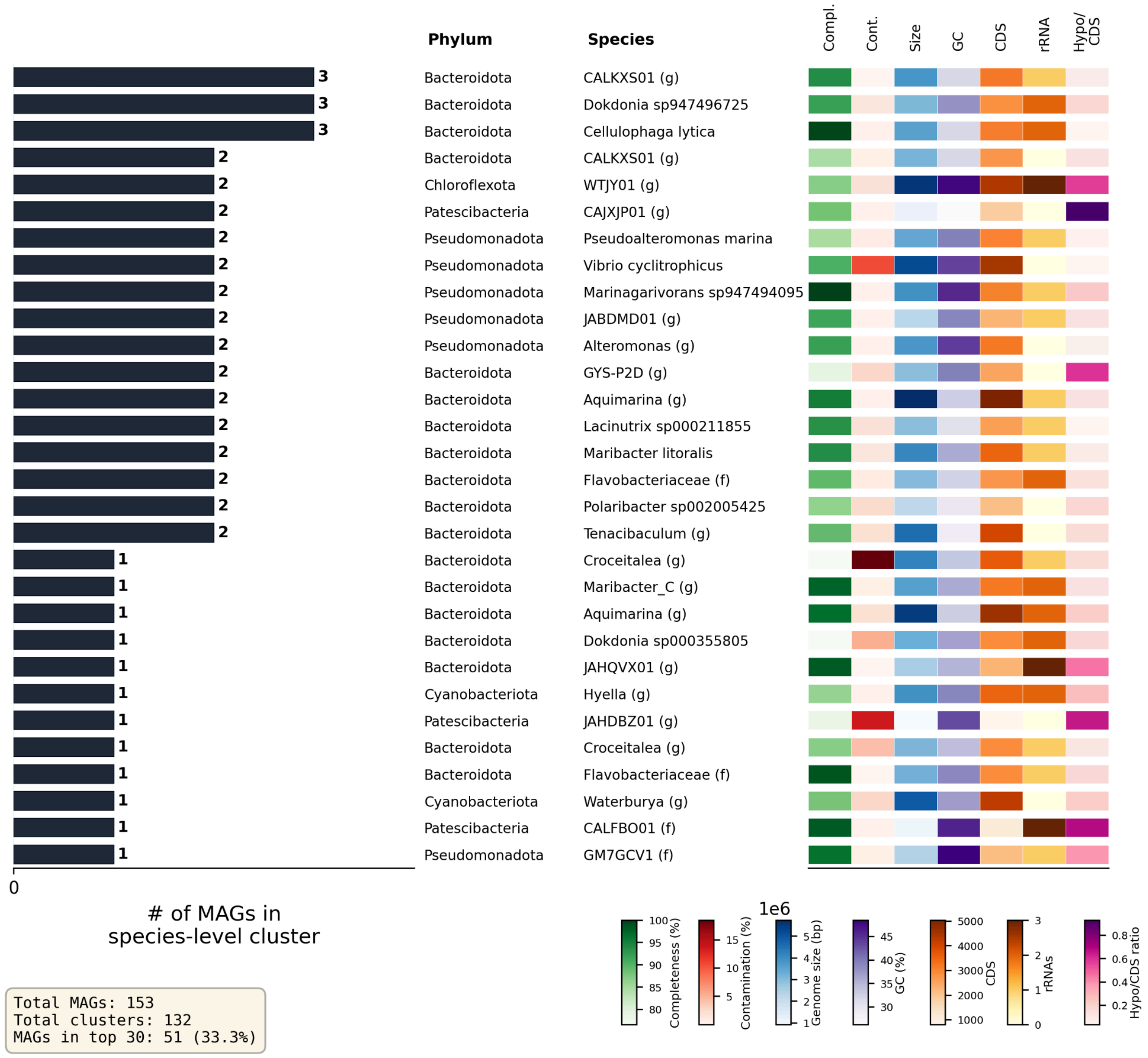


**Supplementary Figure 3. Distribution of top 30 species-level clusters for the macroalgal epiphytic microbiome use case.** The horizontal bar plot shows species-level clusters, as defined by dRep secondary clustering (95% ANI), ranked by the number of MAGs per cluster. Bars represent cluster sizes, with the corresponding numbers of annotated MAGs. Taxonomic assignments (phylum and species) of the representative genomes are shown alongside each cluster. If no species-level assignment was possible, the next highest assigned taxonomic rank is displayed and indicated by its standard abbreviation (e.g., g. = genus). The heatmap panels summarize the quality and genomic characteristics of the representative genomes, including completeness (green), contamination (red), genome size (blue), GC content (purple), number of CDS (orange), number of rRNAs (orange), and the ratio of hypothetical proteins to CDS (pink). Summary statistics, including the total number of MAGs, the total number of species-level clusters, and the proportion of MAGs contained within the largest clusters, are shown in the lower-left corner of the figure.
