## Supplementary Fig. 4 for "FAIRyMAGs - a series of FAIR Galaxy workflows for the generation of metagenome assembled genomes"

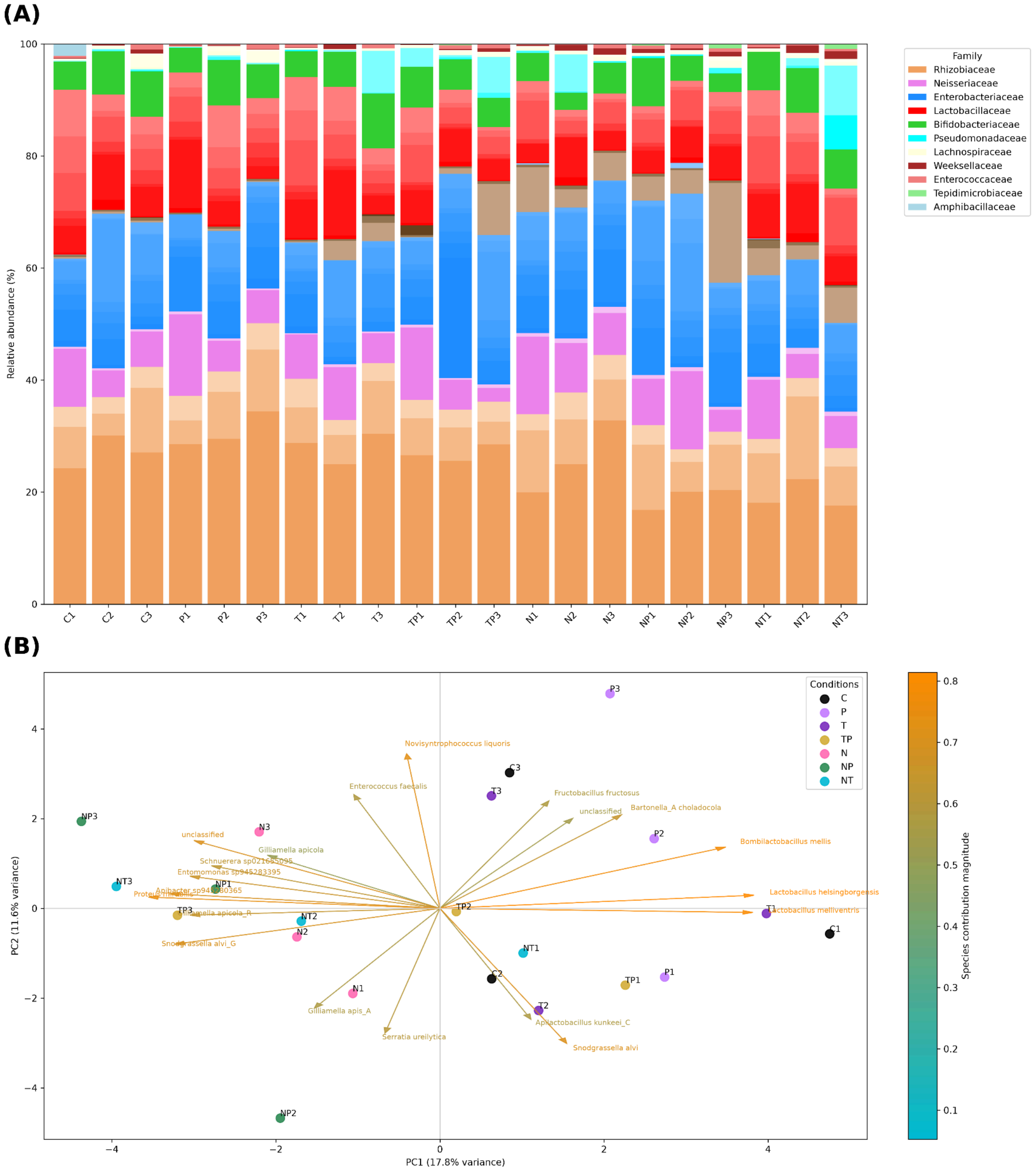


**Supplementary Figure 4. Bee gut bacterial community structure across experimental conditions. (A)** Stacked bar plot of species-level relative abundance of the MQ species-level clusters in individual samples from control (C1–C3), *Pediococcus acidilactici* (P1–P3), thiamethoxam (T1–T3), thiamethoxam plus *P. acidilactici* (TP1–TP3), *Nosema ceranae* (N1–N3), *N. ceranae* plus *P. acidilactici* (NP1–NP3), and *N. ceranae* plus thiamethoxam (NT1–NT3). Colors indicate taxonomic affiliation, with shades differentiating species within families, except for *Proteus spp.* (*P. mirabilis*, light brown), *Providencia* spp. (*Providencia rettgeri* and *Providencia rettgeri_D*, dark brown), and *Arsenophonus apicola*. (black). **(B)** Unsupervised principal component analysis of species-level relative abundance profiles of the MQ species-level clusters across all conditions (n = 3 biological replicates per condition). Sample colors denote treatment groups: control (black), thiamethoxam (purple), *P. acidilactici* (mallow), thiamethoxam plus *P. acidilactici* (gold), *N. ceranae* (pink), *N. ceranae* plus *P. acidilactici* (green), and *N. ceranae* plus thiamethoxam (cyan). Arrows represent the most explanatory species-level variables contributing to the ordination.
